# Fast and unbiased purification of RNA-protein complexes after UV cross-linking

**DOI:** 10.1101/597575

**Authors:** Erika C Urdaneta, Benedikt M Beckmann

**Affiliations:** IRI Life Sciences, Humboldt University, 10115 Berlin, Germany

**Keywords:** RNP, RBP, organic extraction, UV cross-linking

## Abstract

Post-transcriptional regulation of gene expression in cells is facilitated by formation of RNA-protein complexes (RNPs). While many methods to study eukaryotic (m)RNPs rely on purification of polyadenylated RNA, other important regulatory RNA classes or bacterial mRNA could not be investigated at the same depth. To overcome this limitation, we developed Phenol Toluol extraction (PTex), a novel and unbiased method for the purification of UV cross-linked RNPs in living cells. PTex is a fast (2-3 hrs) and simple protocol. The purification principle is solely based on physicochemical properties of cross-linked RNPs, enabling us to interrogate RNA-protein interactions system-wide and beyond poly(A) RNA from a variety of species and source material. Here, we are presenting an introduction of the underlying separation principles and give a detailed discussion of the individual steps as well as incorporation of PTex in high-throughput pipelines.

## 1 Introduction

Cellular gene expression is regulated at different levels. Post-transcriptional regulation comprising mRNA localisation, degradation, translation as well as miRNA-mediated or non-coding RNA-mediated regulation has become a major focus of research in the past years [1, 2]. A hallmark of most eukaryotic mRNA is polyadenylation. Consequently, purification of protein-coding transcripts is facilitated using oligo(dT) beads to enrich for mRNA. In living cells, mRNA is interacting with RNA-binding proteins (RBPs) to form ribonucleoprotein complexes (RNPs), forming a complex network of (m)RNPs in which post-transcriptional regulation is facilitated [3, 4].

Therefore, being able to purify and investigate RNPs is of high importance. In the last years, a set of novel high-throughput techniques has been established in this field. In 2012, the RNA interactome capture (RIC) approach was presented; after UV cross-linking of RBPs to RNA *in vivo*, eukaryotic mRNA is selected using oligo(dT) magnetic beads. The co-purified RBPs are then stringently washed using denaturing conditions and finally identified by mass spectrometry [5, 6]. This resulted in mapping of mRNA-bound proteomes in diverse cell lines and to the identification of hundreds of novel RBPs [2, 7]. However, the RIC approach is limited to eukaryotic mRNA. Investigating other RNA classes (transcripts which are products of RNA polymerase I or III) or mRNA from bacteria and archea cannot be accomplished with this method.

A frequent question is which RNAs are bound by an individual protein. The state-of-the-art method to identify such target RNAs is CLIP (cross-linking and immunoprecipitation) from which many different versions exist now [8]: after UV cross-linking, the RBP of interest is immunoprecipitated using antibodies. The co-purified RNA is then further selected and subsequently sequenced by RNA-Seq.

Both, RIC and CLIP, although been very powerful tools are limited in their scope; being it due to their dependence on poly(A) tails or due to antibody availability. What has been missing is a methodology to purify RNPs in an unbiased fashion. Methods based on RNA *in vivo* labeling with modified nucleotides as RBR-ID [9], RICK [10] and CARIC [11] have been introduced. These approaches utilise modified RNA bases to either identify RBPs or to directly purify RNPs from cells. While these approaches eliminate the focus on polyadenylated RNA, efficient labeling of the biological material emerges as an additional challenge.

Here, we present an approach that separates RNPs solely by physicochemical features that are specific for RNA-protein complexes rather than for individual RNA sequences or for protein epitopes. In our method called Phenol Toluol extraction (PTex) [12]), we are using organic liquid-liquid extractions to enrich UV cross-linked RNPs directly from biological sources such as human cell culture, bacteria or animal tissue.

### 1.1 Theoretical basis

To better understand the PTex approach, we need to first introduce the chemical principles of separating cellular biomolecules by liquid-liquid phase extractions and how to recover RNA and proteins which have been denatured during this procedure. Then, we will discuss the biophysical principles of UV-mediated RNA-protein cross-linking which we exploit for purification of cross-linked RNPs.

#### 1.1.1 The chemistry of extracting nucleic acids using phenol

As starting point for RNP purification, we used extraction of nucleic acids by phenol. This approach has been established already in the 1950s [13] and became a *de facto* standard for RNA isolation when Chomczynski and Sacchi introduced the “single step” method [14] in which isolation of RNAs is achieved by phenolic extractions of cellular lysates using acidic pH and guanidinium thiocyanate.

Phenol extraction exploits the differences in solubility of proteins and nucleic acids in aqueous (polar) and organic (non-polar) solvents. Phenol interacts with hydrophobic amino acid residues of proteins, thus reversing the hydrophobic collapse which is a main driving force of protein folding, and results in denaturation of proteins. In the single step protocol [14], this process is further supported by the chaotropic compound guanidinium thiocyanate. Displaying non-polar/hydrophobic residues, polypeptides are better solvable in phenol (also known as the “like dissolves like” rule) than in water. Nucleic acids on the other hand remain polar, depending on the pH of the solution and dissolve in the aqueous phase. Subsequent centrifugation then separates the two phases; due to its higher density (Table 1), phenol forms the bottom layer and the aqueous phase settles on top. Addition of chloroform or bromochloropropane (BCP) aids in obtaining a sharper phase separation due to their even higher density (Table 1) which reduces carry-over of one of the two phases when pipetting [15].

**Table 1.** Physico-chemical properties of chemicals used for RNA / PTex clRNPs isolation (https://pubchem.ncbi.nlm.nih.gov).

| Property | Chlo-roform | BCP | Phenol | Toluol |
| --- | --- | --- | --- | --- |
| Solubility in water (g/L at 25 °C) | 7.95 | 2.24 | 82.8 | 0.52 |
| Relative density (water= 1) | 1.48 | 1.6 | 1.06 | 0.87 |

During phenolic extraction, RNA and DNA display a different behaviour in respect to their enrichment in the aqueous and interphase at acidic conditions. Nucleobases are primarily in their neutral form at pH 7.2 which is in the physiological (cytosolic) range (Fig. 1), as well as at pH 4.8. Also, the 2’OH group of RNA is in its ionised form at both pH levels. Only the phosphodiester bond has a pK_a_ of 6.0 - 7.0. Thus, the phosphate group of the backbone of DNA and RNA is neutral only at pH 4.8. For DNA, this results in an overall shift from a negatively charged to a neutrally charged molecule. The decrease in polarity of DNA then promotes enrichment in the organic, non-polar phase. RNA however has an additional negative charge due to its 2’OH group with a pK_a_ of 13.0. Additionally and unlike in DNA, the nucleobases are not all paired via H-bonds in a double helix, meaning that unpaired bases can interact with surrounding water molecules, thereby increasing the overall polarity of RNA and its enrichment in the aqueous environment [17]. Subsequent modifications of the method then also allow the recovery of proteins from the organic phase (e.g. see [17, 18]).

**Fig. 1.**
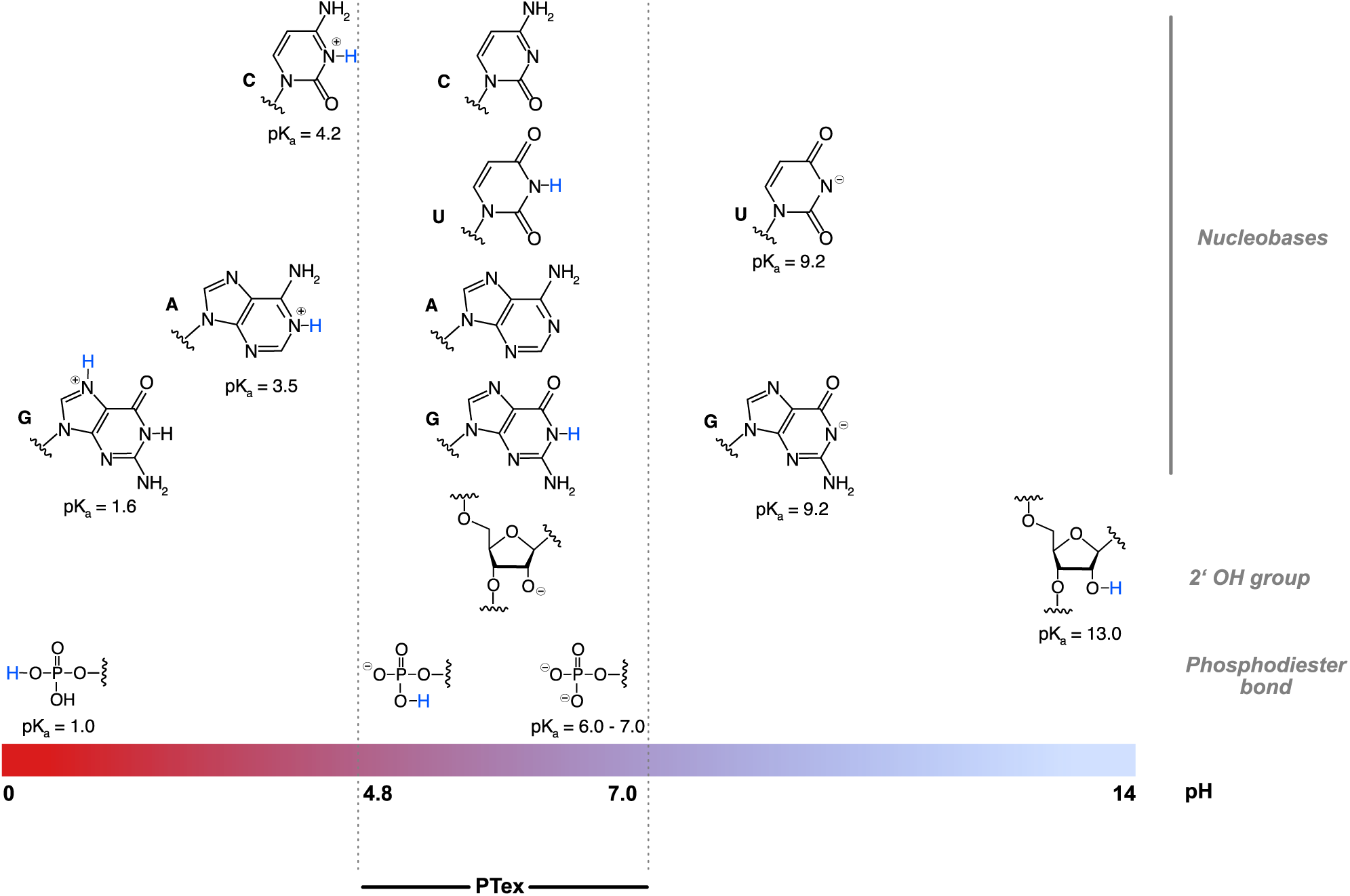
Protonation of nucleotides and PTex. Protonation of sites in RNA and their respective pK_a_s [16]. In the pH range of buffers used in PTex, only the phosphodiester backbone is protonated when shifting from Step1 (pH 7.0) to steps 2/3 (pH 4.8). Nucleobases remain in their neutral form and also the 2’OH of the sugar remains ionised throughout the protocol.

#### 1.1.2 UV cross-linking as a tool to study RNPs

For studying interactions of proteins with RNA, cross-linking of both components of an RNA-protein complex has been used since decades [19, 20]. Particularly useful is the utilisation of short wavelength UV light for cross-linking of RNPs. A major advantage of UV light: it can be applied to living cells (cell culture, tissue) to “capture” RNA-protein interactions *in vivo*, thus preserving physiologically relevant interactions. Before discussing the main advantages and disadvantages however, we aim to introduce the underlying biophysical and chemical events of UV-mediated cross-linking (Fig. 2).

**Fig. 2.**
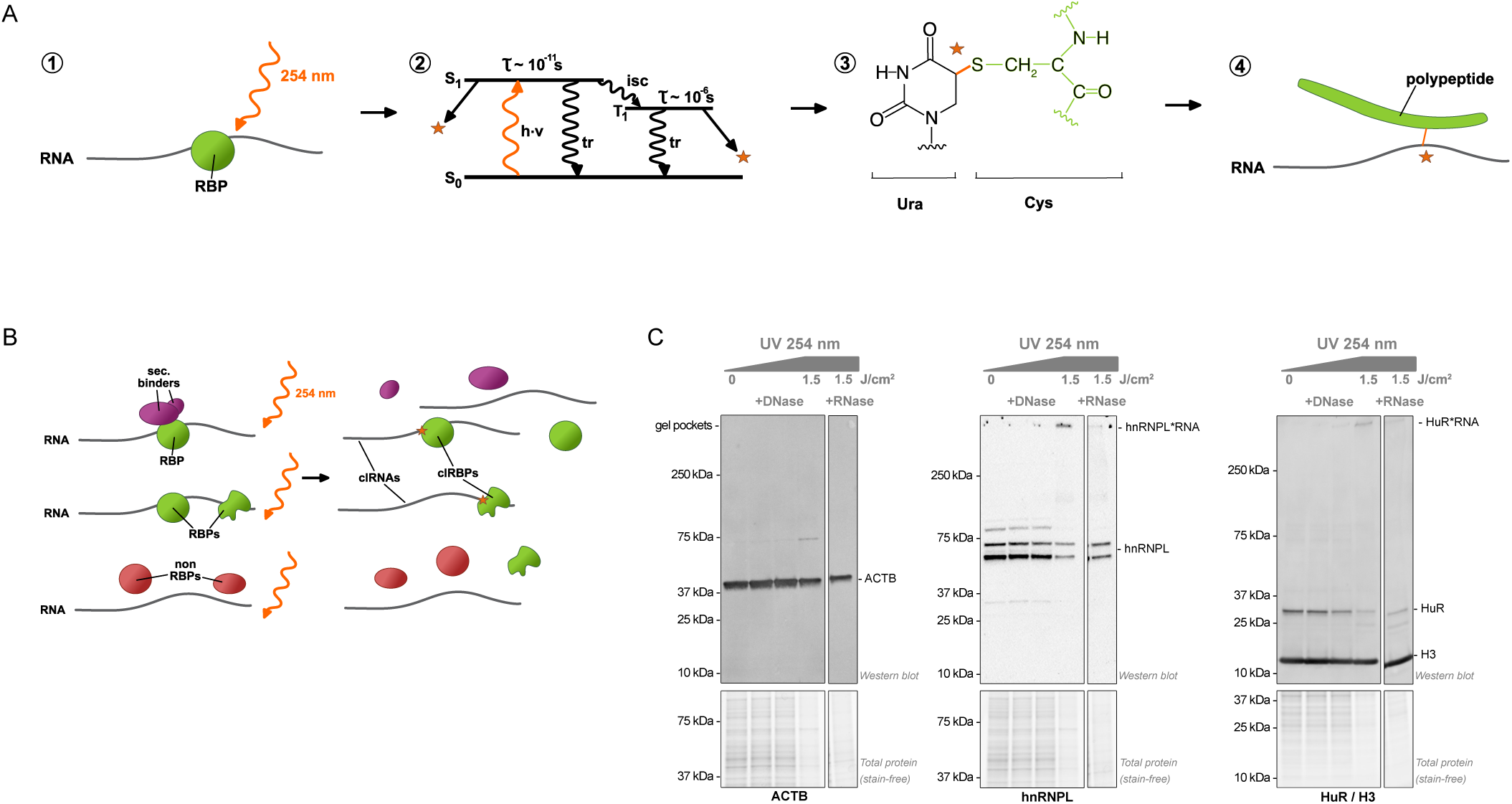
UV cross-linking of RNA-protein complexes. A) Biophysical and chemical basis of UV cross-linking. 1. Irradiation of RNA directly interacting with proteins using low energy short wavelength UV light (254 nm). 2. Simplified Jablonski diagram. Excitation of nucleobases of RNA (h\**ν*) from the ground state S_0_ to an energetically elevated singulet (S_1_) state. Inter state conversion (isc) to a triplet (T_1_) state is possible. The lifetime of S_1_ or T_1_ states are 10 ps and 1 *μ*s, respectively before falling back to the ground state either through thermal relaxation (tr) or by formation of a cross-link to an adjacent amino acid (orange star). Modified from [21]. 3. Example of a cross-link between uracil and cysteine by formation of 5-S-cysteine-6-hydrouracil as determined by [23]. 4. Applying denaturing lysis of successfully cross-linked RNPs results in a hybrid molecule consisting of a nucleic acid and a polypeptide part. B) RBPs (green) can directly interact with RNA in contrast to secondary binders (violet), e.g. protein of RNP complexes interacting solely via protein:protein interactions or non-RBPs (red). UV irradiation at 254 nm can result in covalent cross-links between RBPs and RNA (denoted by an orange star). Note however that UV-induced cross-linking is inefficient and that the majority of the biological sample will remain non-cross-linked. C) Western blots of RBPs from UV-irradiated HEK293 cells (hnRNPL and HuR) demonstrate the low cross-linking efficiency as the main fraction of tested RBPs remains non-cross-linked. Cross-linked HuR and hnRNPL are shifted to a higher molecular mass and stuck in the gel pocket due to the covalently attached RNA, as demonstrated by the reversion of the cross-link signal when cells were treated with Benzonase but not with DNaseI. Note that high UV dosage has adverse effects and results in a general loss of protein (gels in the lower panel), evident also in the case of the non-RBP ACTB detected by Western blot (compare 1.5 J/cm^2^ with lower dosages and in Urdaneta et al. [12]). Histone H3 was used as additional loading control, its relative abundance seems not to be affected by the UV radiation applied.

When irradiating cells with ultraviolet light at 254 nm wavelength, nucleobases of RNA and DNA can absorb the energy of UV light efficiently. Excitation of nucleobases to a higher energetic state S_1_ or T_1_ (when using a low energy UV source), is very short-lived however (Fig. 2A). Within one micro- or even ten picoseconds, the excited state is suspended to the ground state either through thermal relaxation or, if a suitable amino acid is in direct vicinity, by formation of a cross-link. Due to the short time of excitation and since other biological processes such as conformational rearrangement in macromolecules are much slower, cross-links are most likely to form exclusively between components which are in direct contact at the time of irradiation (“zero distance cross-linker”) [19, 21, 22].

What is the actual chemical product of the cross-linking? The first example of a uracil covalently bound to cysteine was reported by Smith and Aplin [23] using NMR spectroscopy and mass spectrometry (Fig. 2A) resulting in formation of 5-S-cysteine-6-hydrouracil. Since then, many more combinations of amino acid/nucleobase cross-links have been investigated (reviewed in [19]). The amino acids cysteine, tyrosine, phenylalanine, arginine, lysine and tryptophane have been reported to be among the most reactive to cross-linking to poly-U [19, 24]. In nucleic acids, pyrimidines are much more efficiently cross-linked in general than purines [25] and with RNA being more reactive than DNA (poly rU > poly rC > poly dT > poly rA) when comparing addition to cysteine [26].

Using UV cross-linking as a starting point for studying RNPs has both, advantages but also disadvantages:

##### Advantages of UV cross-linking

- *In-vivo* RNA-protein interactions can be “frozen” by directly irradiating living cells with UV light.
- Unlike other cross-linkers such as formaldehyde [27], UV light will only cross-link proteins which are in direct contact with (zero distance, see above) but not whole complexes (compare Fig. 2B).
- Photo-irradiation of RNPs results in formation of a covalent bond which is persistent to denaturing conditions; thus permitting to apply stringent conditions during purification

##### Disadvantages of UV cross-linking

- Maybe the most important caveat of UV cross-linking is its very low efficiency. According to own results only up to 5% of a given RBP can be cross-linked to RNA[5, 7, 12, 28]. The efficiency for individual RNA molecules was reported to be higher [29] (note that a single transcript is usually bound by many proteins [3, 4]). Efficiency is even lower when working with tissue or turbid liquid cultures due to the poor penetration of UV light in these media. To overcome this obstacle to some extend, an array of UV bulbs [12, 30] can be used, as e.g. done in a Vari-X-Link device [31].
- Analysis using standard nucleic acid or protein biochemistry techniques can be impaired for cross-linked complexes. The reason is the covalently-attached molecule which adds an additional molecular mass. This often makes it necessary to introduce additional modification steps such as RNase or protease digestion prior to assays like electrophoresis or mass spectrometry. Note however that the additional mass at the cross-linking site can be utilised as a beacon to map RNA-protein interactions at single nucleotide/amino acid resolution [27, 32, 33].
- The formed covalent bond between RNA and protein is thermo-stable and most likely not reversable. Although single studies have suggested reversibility of DNA-protein cross-links by acidic or basic conditions [25], there are no reports that such events occur in RNA-protein covalent bonds to date.

While not being a focus of this paper, note that also RNA-RNA interactions can be cross-linked by UV light. Depending on your biological question, this can be of interest or not [34].

#### 1.1.3 Combining organic extraction and UV cross-linking to investigate RNPs

Having established the effect of UV irradiation on RNPs and knowing the principles of phase separation during phenolic extraction, the behaviour of a cross-linked RNA-protein hybrid (clRNP) in biphasic extractions is of central interest. Displaying physicochemical features of nucleic acid and protein alike, it is reasonable to assume that such clRNPs will accumulate at the phase boundary between aqueous (polar) upper phase and organic (hydropohobic) lower phase. Indeed, previous studies have shown that this area, also known as interphase, contains cross-linked RNPs [20, 35]. However, while these reports used this information to analytically investigate efficiency of UV irradiation, we asked if the differential behaviour of clRNPs in comparison to free RNA and free protein in these extractions could be used for unbiased purification of cross-linked RNPs.

Taken together: UV cross-linking of living cells or tissue will result in a fraction of the cellular RNPs of interest being successfully covalently connected, thereby forming a hybrid molecule with a nucleic acid and a polypetide part. The purpose of PTex is to exploit this two-sided character of the cross-linked molecule using organic liquid-liquid extractions in order to separate them from non-cross-linked molecules.

## 2 Materials and Methods

### 2.1 Cell culture and *in-vivo* cross-linking

Human embrionic kidney cells (HEK293) and HeLa cells (kind gift from Prof. Markus Landthaler (Max-Delbrück Center for Molecular Medicine, Berlin, Germany) and Prof. Andreas Hermann, (Humboldt-Universität zu Berlin, Berlin, Germany)) were grown to 80% confluence at 37°C with 5% CO_2_ on 78 cm^2^ dishes using DMEM high glucose (Dulbecco’s Modified Eagle, glucose 4.5 g/L, Gibco, 41966–029), 10% bovine serum (Gibco, 10270–106), and penicillin/streptomycin (100 U/mL, 0.1 mg/mL; Gibco, 15140–122). Cells in monolayer were washed once with cold phosphate buffer saline (DPBS; Gibco, 10010–015). After removing completely the DPBS, cells (on ice) were irradiated with 0.015, 0.15 and 1.5 J/cm^2^ UV light (λ = 254 nm) in a CL-1000 ultraviolet cross-linker device (Ultra-Violet Products Ltd), collected with cold DPBS, and centrifuged in aliquots of 2-6 x10^6^ cells in 2 mL tubes with safety caps. Cell pellets were stored at −20 or −80 °C (+CL). Non-irradiated cells were used as non cross-link control (-CL).

To test the impact of the different UV dosages (Fig. 2C), HEK293 cells irradiated with 0.015, 0.15 and 1.5 J/cm^2^ UV light (λ = 254 nm) were evaluated by SDS-PAGE and Western blot. HEK293 cell suspensions (1×10^6^/100 μL) were treated with DNAse I (0.2 U/μL, NEB M0303S; NEB DNaseI buffer) or Benzonase (25 U/μL, Merck, 70664; 50 mM Tris, 1 mM MgCL_2_), pH 8.0) during 1 hour at 37 °C. Cells were lysed with 100 μL Laemmli buffer 2x and 20 μL of each sample (including -CL) was loaded in Criterion™ TGX Stain-Free™ Gels (4-20%, BioRad). SDS-PAGE and Western blots were used to reveal the proteins HuR, hnRNPL, ACTB and Histone H3 were performed as described below.

### 2.2 mRNA interactome capture

With the aim of unequivocally track enrichment of clRNPs across phases during the exploratory organic extractions, we prepared mRNA interactome capture (RIC) samples [5] from the *in vivo* cross-linked cells prepared before.

### 2.3 Exploratory organic extractions

We used a simplified screening procedure to probe for the effects of i) phenol:toluol ratios, ii) pH, and iii) native vs. denaturing conditions on their potential to separate molecule classes in liquid-liquid organic extractions:

- RIC samples or whole cells were resuspended in 400 μL of either DPBS (physiological condition) or denaturing solution (solution D: 5.85 M guanidine isothiocyanate (Roth, 0017.3); 31.1 mM sodium citrate (Roth, 3580.3); 25.6 mM N-lauryosyl-sarcosine (PanReac AppliChem, A7402.0100); 1% 2-mercaptoethanol (Sigma), pH 4.8). Then, the aqueous phases were immediately mixed with neutral phenol (400 μL) and 1,3-bromochloropropane (BCP, 100 μL) (Merck, 8.01627.0250); or neutral phenol (Roti-Phenol, Roth 0038.3), toluol (Th.Geyer, 752.1000) and 1,3-bromochloropropane (BCP) (Merck, 8.01627.0250), in a ratio 2:2:1 (500 μL), during 1 min, 2000 r.p.m., 21 °C (Thermomixer, Eppendorf).
- After a short high-speed centrifuging (20.000 xg, 3 min, 4 °C), the upper aqueous phase (aq1) was removed. Then, organic and interphase were mixed with 400 μL of water and 200 μL of ethanol, and centrifuged as before.
- Aqueous-, inter- and organic phases (aq2, inter2, org2) were carefully separated and subjected to ethanol precipitation (9:1, 30 min, −20 °C), followed by centrifuging during 20 min at 20.000 xg, 4 °C. Pellets were dissolved in 30 μL Laemmli buffer. SDS-PAGE and western blotting were performed as described below.

#### 2.3.1 Shifting to aqueous phase assay

We wanted to test whether clRNPs could be shifted from the inter to the aqueous phase: 400 μL of solution D containing 6.6 μg of cross-linked RIC samples from HeLa cells were mixed with 400 μL neutral phenol and BCP (200 μL) during one minute at 2.000 r.p.m, and centrifuged at 20.000 xg, 4 °C for 3 min. After removing 250 μL of the aqueous phase, the inter- and organic phases were mixed with ethanol p.a. (200 μL) and water (400 μL), and centrifuged as before. 3/4 of the aqueous- and organic phases were removed and the resulting interphase mixed (1 min, 2.000 r.p.m, 21 °C) with 400 μL of the indicated buffer (Tables 2, 3, and 200 μL of each, phenol, toluol and BCP, followed by 5 min centrifuging at 20.000 xg, 4 °C. Aqueous and interphase were subjected to ethanol precipitation and prepared for SDS-PAGE.

**Table 2.**
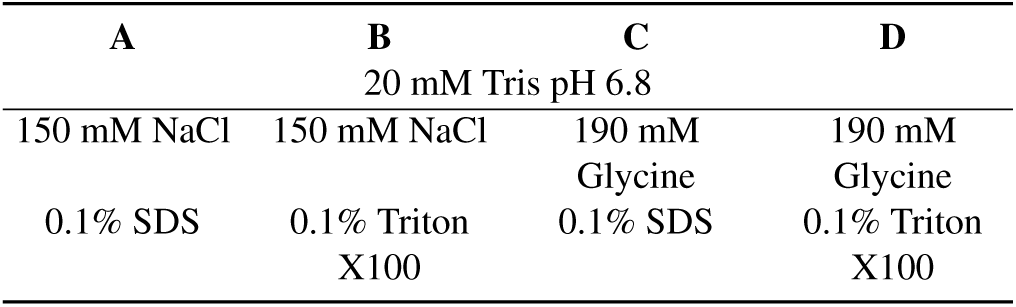
Shifting-to-aqueous phase assay - buffer composition.

**Table 3.**
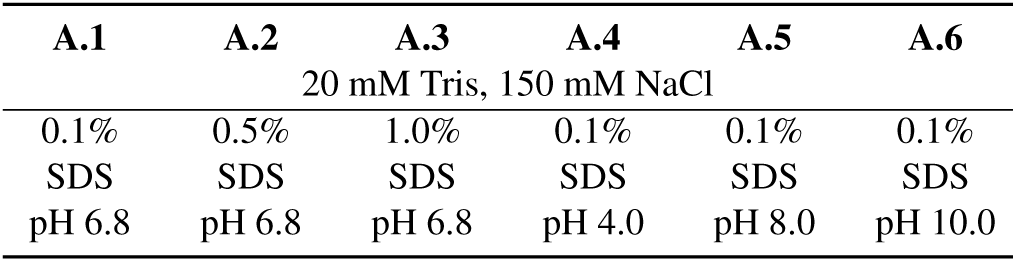
Shifting-to-aqueous phase assay - variations on the pH and detergent concentration of buffer A.

### 2.4 PTex

PTex[12] is a protocol consisting of three fast organic extractions. When extracting 2-8 samples simultaneously, each module (step) can be performed within 10 min; ethanol precipitation and dissolving of the pellet can be completed in about 2 hours. For a convenient reference at the bench, a protocol in form of a flyer is available as Supplementary Information.

- Step 1: HEK293 cell suspensions in 600 μL DPBS (2×10^6^ cells, ±CL) were mixed with 200 μL of each: neutral phenol, toluol and BCP for 1 min (21 °C, 2000 r.p.m, Eppendorf ThermoMixer) and centrifuged 20.000 ×g for 3 min, 4 °C.
- Step 2: The upper aqueous phase (aq1) was carefully removed and transferred to a new 2 mL tube containing 300 μL of solution D. Then, 600 μL neutral phenol and 200 μL BCP were added, mixed and centrifuged as before. After phase separation, the upper three quarters of aq2 and of org2 were removed (in this order, with the help of a syringe with a blunt needle).
- Step 3: The resulting interphase (int2) was kept in the same tube and mixed with 400 μL water, 200 μL ethanol p.a., 400 μL neutral phenol and 200 μL BCP (1 min, 21 ° C, 2000 r.p.m, Eppendorf ThermoMixer) and centrifuged as previously. Three quarters of aq3 and org3 were carefully removed as before, while int3 was precipitated with 9 volumes of ethanol at −20 °C (30 min to overnight), followed by centrifugation at 20.000 xg, 30 min and 4 °C. Pellets were left to dry under the hood for a maximun of 10 min before dissolving.

For PTex step-by-step analysis, all phases were transferred to 5 mL tubes for ethanol precipitation. Pellets dried under the hood for max. 10 min were dissolved with 20 μL Laemmli buffer at 95 °C for 5 min (int1 was dissolved in 100 μL of Laemmli buffer).

### 2.5 Analysis of DNA carry-over during PTex steps

A PTex step-by-step analysis was carried out as described in [12], using 200 ng of pUC19 or gDNA from HEK293 cells. PCRs were applied to amplify a fragment of the genes LacZ (324 nt, forward 5’-AGA GCA GAT TGT ACT GAG-3’and M13-reverse 5’-CAG GAA ACA GCT ATG ACC), or IL3 (574 bp, forward 5’-GAT CGG ATC CTA ATA CGA CTC ACT ATA GGC GAC ATC CAA TCC ATA TCA AGG A-3’ and reverse 5’-GAT CAA GCT TGT TCA GAG TCT AGT TTA TTC TCA CAC-3’). DNA products from each phase were analysed by electrophoresis using 1% agarose gels, Supplementary Figure 12.

### 2.6 Pre-PTex RNase treatment

PTex shown in Fig. 5 was performed using 600 μL HEK293 cells (±CL) previously treated with 2000 U/mL Benzonase (Merck, 70664) in the recommended buffer (50 mM Tris, 1 mM MgCL2, pH 8.0) during 1 h at 37 °C and 1000 rpm (ThermoMixer, Eppendorf), as described in [12]. Following ethanol precipitation, pellets were directly dissolved in 40 μL Laemmli buffer (2×), electrophoresed and blotted as indicated below.

**Fig. 3.**
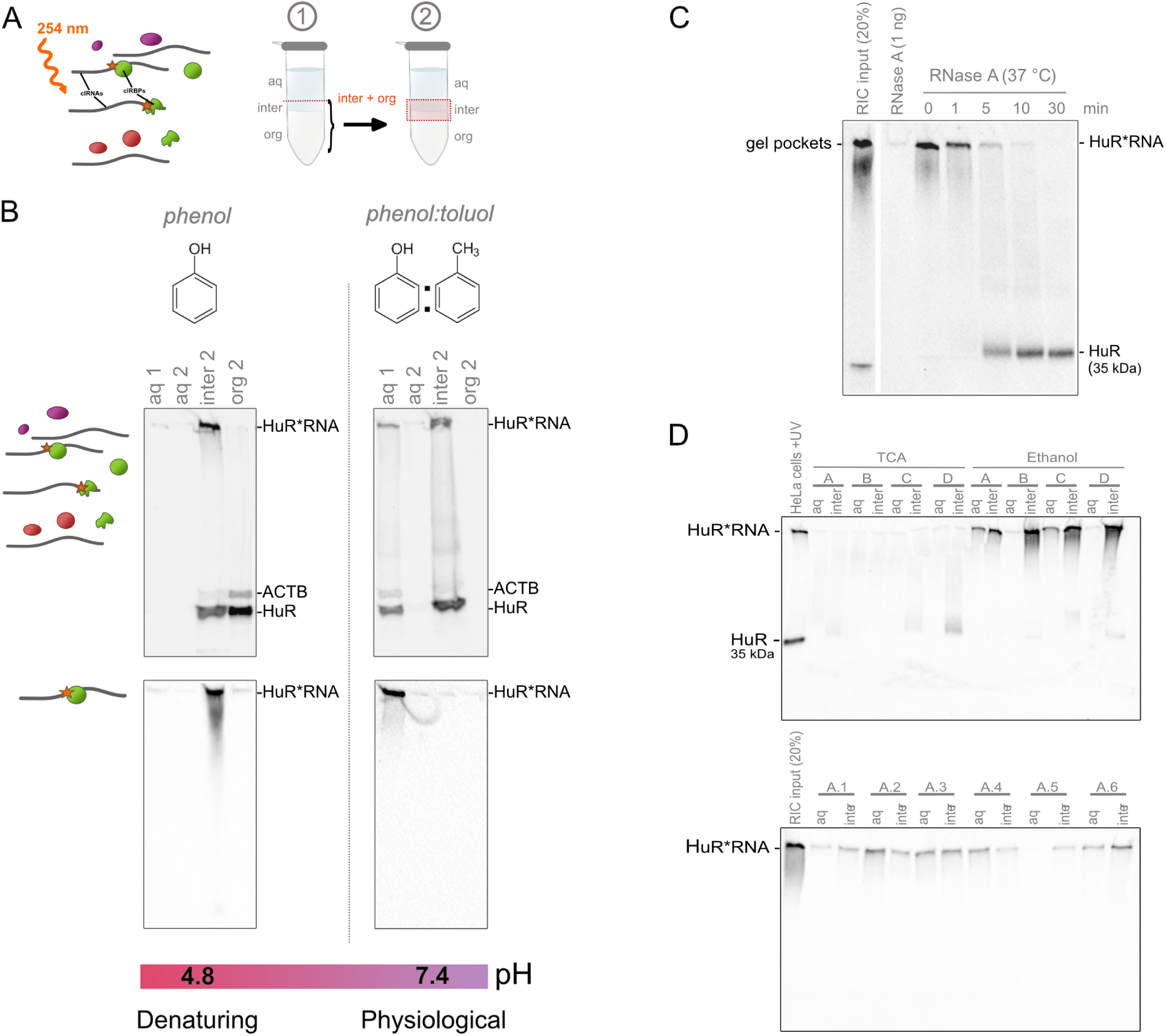
Development of the PTex method. (A) Scheme of the exploratory extractions: UV-irradiated cell pellets resuspended in either denaturing solution D or neutral buffer (DPBS, pH 7.4), mixed with phenol or phenol-toluol, respectively. After centrifuging, the upper aqueous phase was removed and the inter-organic phases mixed and re-extracted with ethanol and water. (B) Phase composition alters the partitioning of clRNPs during organic extractions: (left) in the context of cell lysates (upper panel) or RIC samples (lower panel), phenolic extraction under denaturing conditions at pH 4.8 promotes the migration of the HuR-RNA complexes (HuR*RNA) to the interphase while when a mixture of phenol-toluol and physiological conditions (DPBS, pH 7.4) is applied, the HuR-RNA complexes can be detected in the aqueous phase. (C) HuR-RNA complexes display a reduced electrophoretical mobility (signal at the pockets of the gel) that can be reverse by RNase A treatment (HuR, 35 kDa). (D) RIC-purified complexes subjected to extractions with phenol under denaturing conditions at pH 4.8 can not be fully reversed to the aqueous phase.

**Fig. 4.**
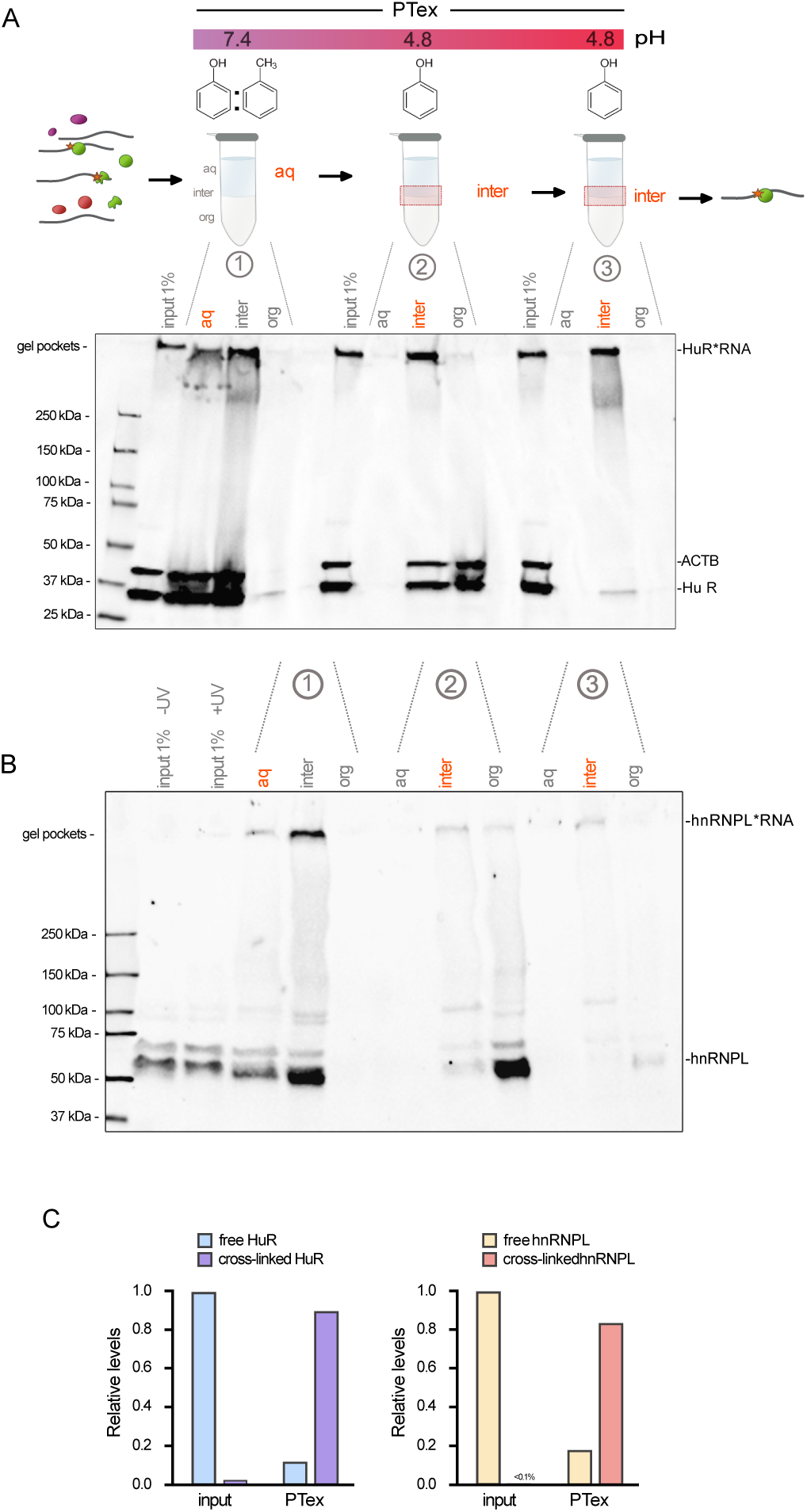
PTex, step-by-step. (A) PTex intermediary steps analysis of HuR and ACTB (3 extractions with 3 phases each). Western blot against HuR (ELAVL1, 35 kDa) shows that UV-cross-linked HuR-RNA complexes (gel pockets) are successfully enriched by PTex whereas the non-RBP ACTB (42 kDa) is completely removed (step 3, interphase) (aqueous phase= aq; interphase= inte; organic phase= org). (B) PTex intermediary steps analysis of hnRNPL (two isoforms: 64.1 and 50.5 kDa). (C) Quantification of (A) and (B): relative enrichment of cross-linked HuR and hn-RNPL by PTex calculated as described in. [12]

**Fig. 5.**
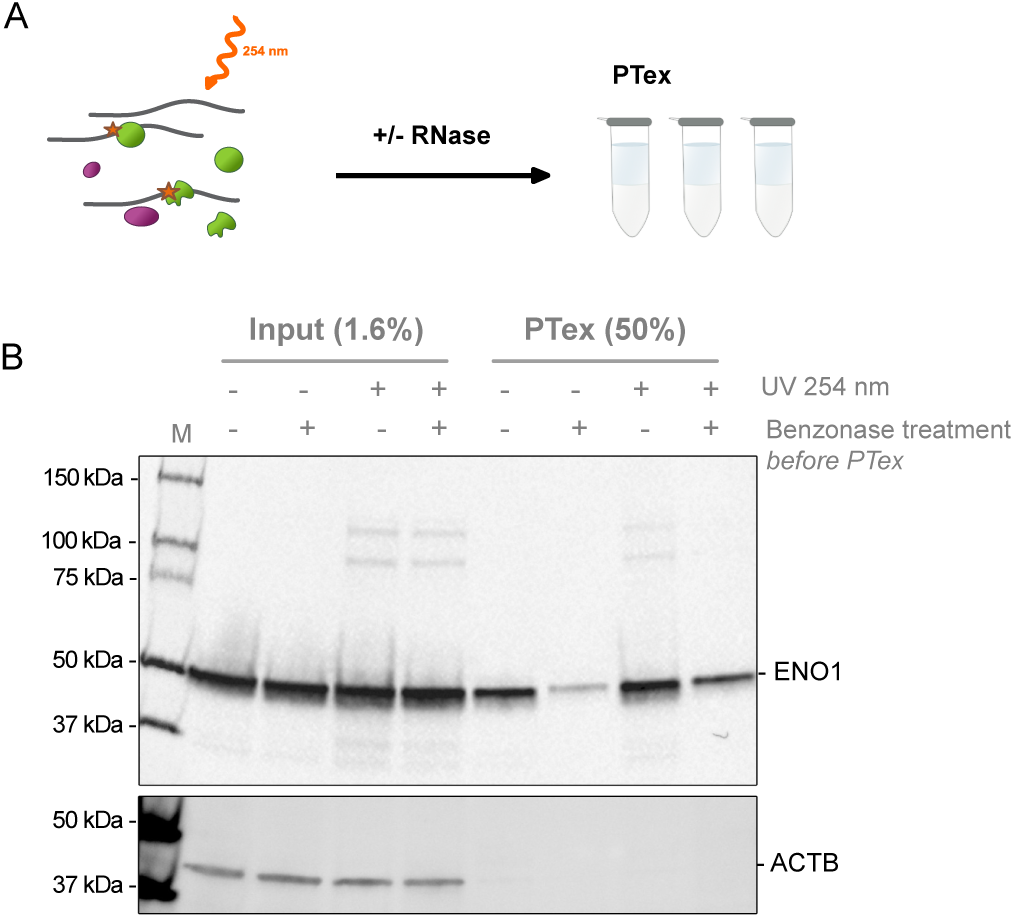
PTex enriches for unconventional RBPs (enigmRBPs [28]) in a RNA-dependent manner. A) HEK293 cells irradiated with 0.25 J/cm^2^ were RNase-treated and subjected to PTex. B) RNase treatment prior to PTex strongly reduced the recovery of ENO-1 when compared with controls (upper blot), while the non-RBP control beta actin (ACTB) was efficiently depleted (lower blot).

### 2.7 Protein precipitation and quantification

HEK293 +CL cell pellets were subjected to the Step 1 of the PTex protocol, the resulting lysates were used as probe for testing three different precipitation methods:

- Ethanol precipitation: samples were mixed with 9 volumes of ethanol p.a., incubated at −20 °C during 30 min and centrifuged 30 min at 20.000 xg and 4 °C. Pellets were washed once with cold ethanol 70% followed for 10 min centrifuging.
- 2-propanol precipitation: samples mixed with 3 volumes of 2-propanol were incubated 10 min at room temperature and centrifuged 20.000 xg, 20 min at 4 °C. Pellets were washed as described above.
- TCA precipitation: cold trichloroacetic acid was added to samples in ratio 0.25:1, followed by 10 min incubation on ice. Samples were centrifuged during 10 min at 20.000 xg, and pellets washed once with cold acetone and centrifuged again.

All supernatants were carefully removed and pellets left to dry under the hood for 10 min. Samples were dissolved with 100 μL of one of the following: water, TE (20 mM Tris, 1 mM EDTA, pH 7.6), or TED buffer (TE supplemented with 0.03% DDM [n-Dodecyl β-D-maltoside]), at 56 °C during 20 min. Samples were spun down at 1000 xg, for 2 min to separate soluble from insoluble material. Protein quantification was determined by measuring the absorbance at λ280 nm in a Nanodrop 2000. Remaining samples were electrophoresed and HuR protein detected by Western blot as detailed below.

### 2.8 Western blotting

Standard techniques were used for western blotting. Samples were electrophoresed on SDS-PAGE gradient gels 4–20% (Criterion™ TGX Stain-Free™ Gels, BioRad), proteins transferred onto nitrocellulose membranes 0.2 μ (Bio-Rad), and membranes blocked during 30 min with PBST-M (10 mM phosphate, 2.7 mM potassium chloride, 137 mM sodium chloride, pH 7.4 0.1% tween 20 (Sigma), 5% milk). Blocked membranes were incubated with 0.1–1.0 μg/mL of the respective antibody overnight at 4°C (or 2 h room temperature). Antibodies targeted the proteins HuR (1:1000, Proteintech, 11910–1-AP), hnRNPL (1:1000, Proteintech 18354-1-AP), ACTB (1:1000, Proteintech, 66009-1-Ig), ENO1 (1:1000, Proteintech 11204-1-AP) and Histone H3 (1:1000, abcam, ab21054). Antibody binding was detected using anti-mouseHRP (1:5000, Proteintech, SA00001-1) or anti-rabbitHRP (1:2500, Proteintech, SA00001-2) and Clarity ECL Western Blotting Substrate for chemiluminescence in a ChemiDocMP imaging system (BioRad).

### 2.9 Electrophoretic mobility shift assay

In order to demonstrate that the accumulation of proteins in the gel’s pockets correspond with the presence of RNA-protein complexes, an electrophoretic mobility shift assay (EMSA) was performed: approximately 15 μg of mRNA interactome capture sample from *in-vivo* cross-linked HeLa cells were subjected to PTex. The resulting pellets were dissolved in 150 μL of ultra-pure water at room temperature, mixed with RNaseA (1 ng) and incubated at 37 °C; aliquots of 20 μL were taken at different time points: 0, 1, 5, 10, and 30 minutes. Aliquots were immediately mixed with 5 μL of 6x Laemmli buffer, heated at 95 °C for 5 min and used for SDS-PAGE and western blotting as described above.

## 3 Results

### 3.1 Establishment of the PTex approach

#### 3.1.1 Phenol-toluol ratios

For a start, we focused on understanding the influence of the different chemical compounds on the clRBPs partitioning during the extraction. For this, we established a base protocol using phenol or phenol-toluol and BCP as organic phase, with either a neutral buffer (PBS, pH 7.4) or the denaturing solution D (pH 4.8) as aqueous phase (Fig. 3A,B). We used HuR (ELAVL1), a well established 35 kDa RNA-binding protein [36] which cross-links efficiently at 254 nm UV light for our setup experiments.

First to notice was an accumulation of HuR in the inter-phases (Fig.3B, upper panel). After UV irradiation, the newly formed RNA-protein hybrid molecules display in a higher molecular weight complex with a reduced mobility when electrophoresed, in this case demonstrated by a signal in the pockets of the gel when detecting the RBP HuR in western blots. This signal is reversed at its normal molecular weight (35 kDa) when digesting the samples with RNase A (Fig. 3C). We used this characteristic as beacon for tracing the migration of the clRNPs between phases during the exploratory extractions: partially lysed cells, membranes and other cellular debris accumulated in the interphase (Supplementary Figure 1), therefore, a major fraction of free or cross-linked HuR (clHuR) were found in this phase which in turn masked the real migration of the clRNPs. Interestingly, two different patterns could be seen: clHuR can be found in the aqueous phase only in the context of extractions using phenol-toluol and neutral buffer. On the other hand, in phenolic extractions under denaturing conditions the majority of the clHuR signal accumulates in the interphase, while unbound HuR could be detected also in the organic phase. More importantly, this findings were corroborated when performing the extractions using pre-purified clRNPs (RIC-samples, Fig. 3B, lower panel).

Due to contamination of the interphase with cellular debris (Supplementary Figure 1), we wanted to investigate whether clRNPs could be shifted from inter- to aqueous phase in a consecutive step. For this we performed extractions using phenol-BCP and solution D, and RIC-purified clRNPs as input material. Then, a third extraction was applied using different buffer compositions as aqueous phase (Fig. 3D; Table 2,3). Although we detected clHuR signal from the aqueous phases, in all cases the complexes partially remained in the interphase.

### 3.2 PTex, step-by-step

To purify clRNPs directly from cells, we combined the previous approaches: taking advantage of the observation that the phenol-toluol mixture under physiological conditions separates soluble RNA, proteins and cross-linked complexes from cellular debris, lipids and the vast majority of DNA, we incorporated this extraction composition as the first step in our protocol [12](Fig. 4A). A phenol-toluol-BCP mixture and neutral buffer (DPBS, pH 7.4) mediated a shift of clRNPs to the aqueous phase separating them from cell contaminants and the vast majority of DNA (Supplementary Figure 1, 12 and [12]). A second extraction with phenol-BCP and a highly denaturing aqueous phase promoted the enrichment of clRNPs in the interphase, while unbound RNAs remained in the aqueous phase and unbound proteins migrated to the organic phase. Finally, a third extraction with phenol-BCP, ethanol and water further removed remaining free proteins. We tested this for several RNA-binding proteins as well as for non-RBPs [12]. Here, HuR, which cross-links very efficiently (up to 1% of the cellular protein can be cross-linked to RNA; Fig. 4 A), and hnRNPL for which UV cross-linking is much less efficient (only up to 0.1% of the cellular protein is cross-linkable (personal communication Oliver Rossbach, University Giessen, Germany; Fig. 4B) are shown to demonstrate two different RBPs that can be investigated by PTex. Considering that in the case of hnRNPL, 99.9% of the protein is not cross-linked (and can thus be considered as contaminant/background in downstream applications), PTex-purified hnRNPL is largely depleted by the free protein and consists of 82% RNA-bound protein (compare input and interphase 3 in Fig. 4 B,C).

In earlier work, we and others found that a surprisingly large number of enzymes of intermediary metabolism bind to RNA *in vivo*, among them the majority of enzymes in glycolysis [28, 37]. For enolase (ENO-1), theses findings were confirmed and target RNAs were identified recently [38]. Here, we show that using PTex, it is likewise possible to extract RNA-binding enzymes such as ENO-1 (Fig. 5). In this experimental setup, we also used RNase treatment after *in vivo* UV irradiation but before proceeding with the PTex protocol. This was done to demonstrate that efficient recovery of ENO-1 by PTex relies not only on UV cross-linking, but also on presence of covalently bound RNA.

Worth to mention is the use of phase-lock gel tubes for a sharper phase separation during the extraction: although this system proved to be useful for RNA phenol-based extractions where the aqueous -RNA containg-phase is well separated from the organic solution, during PTex and related methods [12, 39, 40], clRNPs accumulate in the interphase which is a transitioning space between the lower end of the aqueous phase and the upper part of the organic phase. When the gel in the phase-lock tube separates the aqueous and organic phases, the interphase does not form any longer, and by consequence the clRNPs get misplaced (probably embedded within the gel; data not shown).

As described in our previous work [12], PTex recovers around 30% of clRNPs. However, is this discrete yield solely due to the extraction procedure or are the protein precipitation and dissolving steps also affecting the recovery of the complexes? To answer this question we submitted fractions of the same cell lysate to three of the most commonly used protein/nucleic-acids precipitation methods: ethanol, 2-propanol and trichloroacetic acid (TCA), then resuspending the pellets in either water, TE or TED buffers (see section 2.5). Even though the different precipitation methods rendered an overall similar protein recovery, TCA precipitation produced a rather insoluble pellet (Supplementary Figures 2,3). Besides the low protein recovery obtained by the methods tested, alcoholic precipitations resulted in a sample that dissolved easier and preserved RNA better (Fig.3D, Supplementary Figures 2,3). Nonetheless, particular attention on the precipitation method and dissolving strategy must be taken when preparing PTex samples for mass spectrometry as RNA and traces of the reagents could compromise the integrity of the LC columns (see [12] for detailed information on sample preparation for mass spectrometry).

## 4 Discussion

As demonstrated for hnRNPL, only ∼1 out of 1000 molecules can be cross-linked to RNA *in vivo*. Each down-stream application faces the challenge to remove 99.9% non cross-linked background protein. Similar problems arise when performing RNA interactome capture approaches: the majority of binding sites in poly(A) RNA will not contain cross-linked RBP of interest. PTex is an approach to overcome those obstacles and is based on two principles: i) separate RNA and proteins from other cellular molecules (step 1) and ii) remove free RNA and free proteins from clRNPs in an unbiased fashion (step 2 & 3). Simultaneously to our method, two similar approaches to purify clRNPs employing phenolic extractions have been published (XRNAX [39], OOPS [40]).

### 4.1 Applications

We designed PTex as a versatile method. Fig. 3C is of particular importance in this respect: here, we are using PTex-purified, *in vivo* cross-linked HuR-RNA complexes for RNase digestion to demonstrate that the signal in the gel pocket was indeed a result of UV-mediated covalent cross-linking of RNA and proteins. More importantly however is the observation that the HuR clRNP can be subjected to enzymatic digestion of RNA after PTex, meaning that our approach does not only remove the majority of free RNA and proteins but leaves the enriched RNPs amenable to downstream applications. We obtained similar results when using Proteinase K for proteolytic digestion (data not shown). Hence, PTex can be utilised in a modular fashion and incorporated into complex workflows. A plethora of potential applications come to mind, e.g. to reduce the amount of necessary antibody for a RBP in CLIP-type experiments. This renders PTex particularly interesting for high-throughput approaches or when conducting serial experiments. In this light, we already used PTex to simplify a PAR-CLIP [32] workflow by replacing extraction of radio-labeled RNPs from gels/membranes by phenolic extraction [12].

Another challenging aspect of UV cross-linking is to obtain RNPs from non-cell culture source material. The already very low efficiency of UV irradiation is further decreased when probing tissue samples or liquid cultures such as yeast [28, 37, 41]. We used PTex to directly purify cross-linked HuR from mouse brain samples [12]. This is of particular interest since *in vivo* RNA labeling of whole animals as well as some unicellular species has not been efficiently conducted. The latter however is a prerequisite for other RNP purification techniques such as PAR-CLIP [32], RBR-ID [9], RICK [10] or CARIC [11].

We suggest that PTex has a large potential in RNA biology high-throughput experiments. Using RNA interactome capture, the mRNA-bound proteomes of diverse species and cell types has been determined [2, 7]. Since PTex is not restricted to poly(A) RNA, we used our approach to determine the complete RNA-bound proteome of a human cell: using PTex-purified samples from whole HEK293 cells, we analysed the protein fraction of the clRNPs by mass spectrometry, following an analysis pipeline which we had established before [41]. This allowed us to largely increase the number of RNA-associated proteins [12]. We identified protein groups which were not associated with RNA interaction so far, such as e.g. AAA ATPases. Similar features were found by the other two methods employing phenolic extractions which investigated additional cell types and cellular compartments [39, 40].

Along the same lines: the landscape of bacterial RBPs has not been determined at the same depth as for eukaryotic species; a major issue being the lack of poly(A) RNA necessary for interactome capture [5]. PTex allowed us for the first time to unbiasedly screen for proteins cross-linked to RNA in *Salmonella* Typhimurium [12] while RBPs in *E. coli* were purified by OOPS [40] and TRAPP [38].

### 4.2 Considerations before using UV cross-linking

As mentioned before, UV cross-linking is highly inefficient. Despite that, it has been widely used in studying RNA-protein complexes. However, before using UV cross-linking as a starting point for investigating RNPs, other potential issues should be taken into consideration as well.

First, like RNA, DNA can be cross-linked to proteins by UV light at 254 nm wavelength (see [19] for a comprehensive review on this topic). This makes it even more important, that applications such as PTex select RNA-binding proteins and not DNA-binders alike. Free DNA is depleted during the PTex procedure (see Supplementary Fig. 12) and abundant DNA-binders such as histone H3 and many transcription factors are not captured by PTex [12]. However, a variety of DNA-binders have been reported to also display RNA-binding activity [42]. For these proteins, to distinguish between both activities poses a challenge when using UV irradiation as experimental approach.

Second, there is evidence that UV irradiation can lead to protein-protein cross-links as well [21, 43, 44]. Especially when using UV-based approaches such as RIC or PTex for screening purposes, this could jeopardise the specificity of RBP discovery as secondary, non-direct RNA interactors would be cross-linked to *bona fide* RBPs as well. However, to the best of our knowledge, protein-protein cross-links were exclusively found when using high energy lasers as UV source [21, 22, 43, 44]. RIC, CLIP or PTex experiments on the other hand have so far been conducted using UV bulbs in a Stratalinker (or similar) device in which a complete area is irradiated. Additionally, these protocols classically use energies between 0.2 and 0.4 J/cm^2^_254 nm_ while the studies using a laser applied 1 J_254 nm_ or more to a small area [43, 44]. We and others have used higher UV intensities for RBP discovery as well [12, 38]. However, while we are not aware of a systematic comparison of the UV bulbs we used and UV lasers, Budowsky et al. [21] noted that laser-irradiation results in an additional excitation state H_T_ which cannot be reached when using low energy sources (see Fig. 2A). These findings suggest that the two UV irradiation strategies are different in respect to efficacy of cross-linking in cells, and it remains questionable if the applied energies are comparable at all, e.g. due to differences in penetration of biological material.

Finally, using high UV dosages (we tested up to 1.5 J/cm^2^_254 nm_ from UV bulbs; see Fig. 2C and [12]), we found that prolonged exposure to UV light has adverse effects on RBP recovery. As also shown in Fig. 2C, overall protein levels decrease during the treatment. The exact cause for this is unclear. We are speculating that the UV exposure triggers protease digestion in the living cells. We recommend to avoid such high UV dosages (and/or long exposure times) to keep a cellular state as close as possible to physiological conditions.

### 4.3 Limitations of PTex

We thoroughly tested PTex efficiency before to determine its limitations [12]:

- Enrichment vs. recovery: While largely enriching for cross-linked over free RNP components, clRNP recovery is not complete as some material is lost during the protocol. We have estimated that PTex recovers 25-30% of the initially cross-linked RNPs. If loss of clRNPs (in absolute terms) is not acceptable (e.g. due to scarcity of starting material), applying PTex might impair overall purification success.
- We found that RNA as short as 30 nt could be efficiently purified using PTex when bound to a single RBP. When investigating complexes containing shorter RNA species such as mature miRNA [45, 46], PTex is not the method of choice. Furthermore, we did not systematically compare different protein masses and RNA lengths for purification efficiency by PTex. Minimal RNA length or protein size might differ for other complex compositions.
- During setup of PTex, we noticed that the extraction tube can become saturated, resulting in impaired separation and purification. In this light, a sufficient volume of the PTex reagents in relation to the sample size is important.
- PTex suffers from a technical issue inherited from the single step protocol: removing of individual phases and separating aqueous, inter- and organic phase using a syringe or pipet tip is technically challenging and will to some extend depend on the skill of the experimenter.
- Finally, we find it important to point to a semantic issue. As discussed before [41], proteins cross-linked to RNA are not automatically RNA-binding proteins in the classical sense; the term “RBP” has historically been used for proteins with a role in RNA biology such as RNases, helicases, etc. However, UV-induced covalent bonds will be formed because of physical proximity in the cell and not because of the physiological role of a protein. Structural elements of RNP complexes can easily be cross-linked (and hence purified by PTex) to RNA. Likewise, proteins without RNA-binding activity can be bound RNA [47, 48]. We usually refer to the PTex-purified proteins as “RNA associated” to avoid over-interpretation and confusion with classical RNA-regulating proteins.

## Supporting information

PTex Flyer

## ACKNOWLEDGEMENTS

The authors wish to thank Julie Bohl for technical assistance. Work in the Beckmann lab is supported by the German Research Foundation (DFG; IRTG 2290 and ZUK 75/1 Project 0190-854599). ECU is supported by the Joachim Herz Foundation.

## AUTHOR CONTRIBUTIONS

E.C.U. conducted the experiments. E.C.U. and B.M.B wrote the manuscript.

## Bibliography

1. Sarah F. Mitchell and Roy Parker. Principles and properties of eukaryotic mRNPs. Molecular Cell, 54(4):547–558, May 2014. ISSN 1097-4164. doi:10.1016/j.molcel.2014.04.033.

2. Matthias W. Hentze, Alfredo Castello, Thomas Schwarzl, and Thomas Preiss. A brave new world of RNA-binding proteins. Nature Reviews. Molecular Cell Biology, 19(5):327–341, May 2018. ISSN 1471-0080. doi:10.1038/nrm.2017.130.

3. Niels H. Gehring, Elmar Wahle, and Utz Fischer. Deciphering the mRNP Code: RNA-Bound Determinants of Post-Transcriptional Gene Regulation. Trends in Biochemical Sciences, 42 (5):369–382, May 2017. ISSN 0968-0004. doi:10.1016/j.tibs.2017.02.004.

4. Daniel J. Hogan, Daniel P. Riordan, André P. Gerber, Daniel Herschlag, and Patrick O. Brown. Diverse RNA-binding proteins interact with functionally related sets of RNAs, suggesting an extensive regulatory system. PLoS Biology, 6(10):e255, Oct 2008. ISSN 1545-7885. doi:10.1371/journal.pbio.0060255.

5. Alfredo Castello, Bernd Fischer, Katrin Eichelbaum, Rastislav Horos, Benedikt M. Beckmann, Claudia Strein, Norman E. Davey, David T. Humphreys, Thomas Preiss, Lars M. Steinmetz, Jeroen Krijgsveld, and Matthias W. Hentze. Insights into RNA Biology from an Atlas of Mammalian mRNA-Binding Proteins. Cell, 149(6):1393–1406, jun 2012. ISSN 1097-4172. doi:10.1016/j.cell.2012.04.031.

6. Alexander G. Baltz, Mathias Munschauer, Björn Schwanhäusser, Alexandra Vasile, Ya-suhiro Murakawa, Markus Schueler, Noah Youngs, Duncan Penfold-Brown, Kevin Drew, Miha Milek, Emanuel Wyler, Richard Bonneau, Matthias Selbach, Christoph Dieterich, and Markus Landthaler. The mRNA-bound proteome and its global occupancy profile on protein-coding transcripts. Molecular Cell, 46(5):674–690, Jun 2012. ISSN 1097-4164. doi:10.1016/j.molcel.2012.05.021.

7. Benedikt M. Beckmann, Alfredo Castello, and Jan Medenbach. The expanding universe of ribonucleoproteins: of novel RNA-binding proteins and unconventional interactions. Pflugers Archiv: European journal of physiology, 468(6):1029–1040, Jun 2016. ISSN 1432-2013. doi:10.1007/s00424-016-1819-4.

8. Flora C. Y. Lee and Jernej Ule. Advances in CLIP Technologies for Studies of Protein-RNA Interactions. Molecular Cell, 69(3):354–369, Feb 2018. ISSN 1097-4164. doi:10.1016/j.molcel.2018.01.005.

9. Chongsheng He, Simone Sidoli, Robert Warneford-Thomson, Deirdre C. Tatomer, Jeremy E. Wilusz, Benjamin A. Garcia, and Roberto Bonasio. High-Resolution Mapping of RNA-Binding Regions in the Nuclear Proteome of Embryonic Stem Cells. Molecular Cell, 64(2):416–430, Oct 2016. ISSN 1097-4164. doi:10.1016/j.molcel.2016.09.034.

10. Xichen Bao, Xiangpeng Guo, Menghui Yin, Muqddas Tariq, Yiwei Lai, Shahzina Kanwal, Jiajian Zhou, Na Li, Yuan Lv, Carlos Pulido-Quetglas, Xiwei Wang, Lu Ji, Muhammad J. Khan, Xihua Zhu, Zhiwei Luo, Changwei Shao, Do-Hwan Lim, Xiao Liu, Nan Li, Wei Wang, Minghui He, Yu-Lin Liu, Carl Ward, Tong Wang, Gong Zhang, Dongye Wang, Jianhua Yang, Yiwen Chen, Chaolin Zhang, Ralf Jauch, Yun-Gui Yang, Yangming Wang, Baoming Qin, Minna-Liisa Anko, Andrew P. Hutchins, Hao Sun, Huating Wang, Xiang-Dong Fu, Biliang Zhang, and Miguel A. Esteban. Capturing the interactome of newly transcribed RNA. Nature Methods, 15(3):213–220, Mar 2018. ISSN 1548-7105. doi:10.1038/nmeth.4595.

11. Rongbing Huang, Mengting Han, Liying Meng, and Xing Chen. Transcriptome-wide discovery of coding and noncoding RNA-binding proteins. Proceedings of the National Academy of Sciences of the United States of America, 115(17):E3879–E3887, Apr 2018. ISSN 1091-6490. doi:10.1073/pnas.1718406115.

12. Erika C Urdaneta, Carlos H Vieira-Vieira, Timon Hick, Hans-Herrmann Wessels, Davide Figini, Rebecca Moschall, Jan Medenbach, Uwe Ohler, Sander Granneman, Matthias Selbach, and Benedikt M Beckmann. Purification of cross-linked RNA-protein complexes by phenol-toluol extraction. Nature Communications, 10(1):990, 2019. ISSN 2041-1723. doi:10.1038/s41467-019-08942-3.

13. K. S. Kirby. A new method for the isolation of ribonucleic acids from mammalian tissues. The Biochemical Journal, 64(3):405–408, Nov 1956. ISSN 0264-6021.

14. P. Chomczynski and N. Sacchi. Single-step method of RNA isolation by acid guanidinium thiocyanate-phenol-chloroform extraction. Analytical Biochemistry, 162(1):156–159, Apr 1987. ISSN 0003-2697. doi:10.1006/abio.1987.9999.

15. P. Chomczynski and K. Mackey. Substitution of chloroform by bromo-chloropropane in the single-step method of RNA isolation. Analytical Biochemistry, 225(1):163–164, Feb 1995. ISSN 0003-2697. doi:10.1006/abio.1995.1126.

16. Pallavi Thaplyal and Philip C. Bevilacqua. Experimental approaches for measuring pKa’s in RNA and DNA. Methods in Enzymology, 549:189–219, 2014. ISSN 1557-7988. doi:10.1016/B978-0-12-801122-5.00009-X.

17. P Zumbo. Phenol-chloroform Extraction.

18. Soroth Chey, Claudia Claus, and Uwe Gerd Liebert. Improved method for simultaneous isolation of proteins and nucleic acids. Analytical Biochemistry, 411(1):164–166, Apr 2011. ISSN 1096-0309. doi:10.1016/j.ab.2010.11.020.

19. M. D. Shetlar. Cross-Linking of Proteins to Nucleic Acids by Ultraviolet Light. Photochem. Photobiol. Rev., (5):105–197, 1980.

20. A. Favre, G. Moreno, M. O. Blondel, J. Kliber, F. Vinzens, and C. Salet. 4-Thiouridine photosensitized RNA-protein crosslinking in mammalian cells. Biochemical and Biophysical Research Communications, 141(2):847–854, Dec 1986. ISSN 0006-291X.

21. E. I. Budowsky, M. S. Axentyeva, G. G. Abdurashidova, N. A. Simukova, and L. B. Rubin. Induction of polynucleotide-protein cross-linkages by ultraviolet irradiation. Peculiarities of the high-intensity laser pulse irradiation. European Journal of Biochemistry, 159(1):95–101, Aug 1986. ISSN 0014-2956.

22. I. G. Pashev, S. I. Dimitrov, and D. Angelov. Crosslinking proteins to nucleic acids by ultra-violet laser irradiation. Trends in Biochemical Sciences, 16(9):323–326, Sep 1991. ISSN 0968-0004.

23. K. C. Smith and R. T. Aplin. A mixed photoproduct of uracil and cysteine (5-S-cysteine-6-hydrouracil). A possible model for the in vivo cross-linking of deoxyribonucleic acid and protein by ultraviolet light. Biochemistry, 5(6):2125–2130, Jun 1966. ISSN 0006-2960.

24. M. D. Shetlar, J. Carbone, E. Steady, and K. Hom. Photochemical addition of amino acids and peptides to polyuridylic acid. Photochemistry and Photobiology, 39(2):141–144, Feb 1984. ISSN 0031-8655.

25. P. R. Paradiso, Y. Nakashima, and W. Konigsberg. Photochemical cross-linking of protein. Nucleic acid complexes. The attachment of the fd gene 5 protein to fd DNA. The Journal of Biological Chemistry, 254(11):4739–4744, Jun 1979. ISSN 0021-9258.

26. K. C. Smith and D. H. Meun. Kinetics of the photochemical addition of [35S] cysteine to polynucleotides and nucleic acids. Biochemistry, 7(3):1033–1037, Mar 1968. ISSN 0006-2960.

27. Amol Panhale, Florian M. Richter, Fidel Ramírez, Maria Shvedunova, Thomas Manke, Gerhard Mittler, and Asifa Akhtar. CAPRI enables comparison of evolutionarily conserved RNA interacting regions. Nature communications, 10(1):2682, Jun 2019. ISSN 2041-1723. doi:10.1038/s41467-019-10585-3.

28. Benedikt M. Beckmann, Rastislav Horos, Bernd Fischer, Alfredo Castello, Katrin Eichelbaum, Anne-Marie Alleaume, Thomas Schwarzl, Tomaz Curk, Sophia Foehr, Wolfgang Hu- ber, Jeroen Krijgsveld, and Matthias W. Hentze. The rna-binding proteomes from yeast to man harbour conserved enigmrbps. Nature Communications, 6:10127, Dec 2015. ISSN 2041-1723. doi:10.1038/ncomms10127.

29. A. J. Wagenmakers, R. J. Reinders, and W. J. van Venrooij. Cross-linking of mRNA to proteins by irradiation of intact cells with ultraviolet light. European Journal of Biochemistry, 112(2):323–330, Nov 1980. ISSN 0014-2956.

30. Rob van Nues, Gabriele Schweikert, Erica de Leau, Alina Selega, Andrew Langford, Ryan Franklin, Ira Iosub, Peter Wadsworth, Guido Sanguinetti, and Sander Granneman. Kinetic CRAC uncovers a role for Nab3 in determining gene expression profiles during stress. Nature Communications, 8(1):12, Apr 2017. ISSN 2041-1723. doi:10.1038/s41467-017-00025-5.

31. Vari-X-Link Homepage. https://www.vari-x-link.com.

32. Markus Hafner, Markus Landthaler, Lukas Burger, Mohsen Khorshid, Jean Hausser, Philipp Berninger, Andrea Rothballer, Manuel Ascano, Anna-Carina Jungkamp, Mathias Munschauer, Alexander Ulrich, Greg S. Wardle, Scott Dewell, Mihaela Zavolan, and Thomas Tuschl. Transcriptome-wide identification of rna-binding protein and microrna target sites by par-clip. Cell, 141(1):129–141, Apr 2010. ISSN 1097-4172. doi:10.1016/j.cell.2010.03.009.

33. Katharina Kramer, Timo Sachsenberg, Benedikt M. Beckmann, Saadia Qamar, Kum-Loong Boon, Matthias W. Hentze, Oliver Kohlbacher, and Henning Urlaub. Photo-cross-linking and high-resolution mass spectrometry for assignment of rna-binding sites in rna-binding proteins. Nature Methods, 11(10):1064–1070, Oct 2014. ISSN 1548-7105. doi:10.1038/nmeth.3092.

34. Stefanie Grosswendt, Andrei Filipchyk, Mark Manzano, Filippos Klironomos, Marcel Schilling, Margareta Herzog, Eva Gottwein, and Nikolaus Rajewsky. Unambiguous identification of miRNA:target site interactions by different types of ligation reactions. Molecular Cell, 54(6):1042–1054, Jun 2014. ISSN 1097-4164. doi:10.1016/j.molcel.2014.03.049.

35. R. Brimacombe, W. Stiege, A. Kyriatsoulis, and P. Maly. Intra-RNA and RNA-protein cross-linking techniques in Escherichia coli ribosomes. Methods in Enzymology, 164:287–309, 1988. ISSN 0076-6879.

36. Svetlana Lebedeva, Marvin Jens, Kathrin Theil, Björn Schwanhäusser, Matthias Selbach, Markus Landthaler, and Nikolaus Rajewsky. Transcriptome-wide analysis of regulatory interactions of the RNA-binding protein HuR. Molecular Cell, 43(3):340–352, Aug 2011. ISSN 1097-4164. doi:10.1016/j.molcel.2011.06.008.

37. Ana M. Matia-González, Emma E. Laing, and André P. Gerber. Conserved mRNA-binding proteomes in eukaryotic organisms. Nature Structural & Molecular Biology, 22(12):1027–1033, Dec 2015. ISSN 1545-9985. doi:10.1038/nsmb.3128.

38. Vadim Shchepachev, Stefan Bresson, Christos Spanos, Elisabeth Petfalski, Lutz Fischer, Juri Rappsilber, and David Tollervey. Defining the RNA interactome by total RNA-associated protein purification. Molecular systems biology, 15(4):e8689, Apr 2019. ISSN 1744-4292. doi:10.15252/msb.20188689.

39. Jakob Trendel, Thomas Schwarzl, Rastislav Horos, Ananth Prakash, Alex Bateman, Matthias W. Hentze, and Jeroen Krijgsveld. The Human RNA-Binding Proteome and Its Dynamics during Translational Arrest. Cell, 176(1-2):391–403.e19, Jan 2019. ISSN 1097-4172. doi:10.1016/j.cell.2018.11.004.

40. Rayner M. L. Queiroz, Tom Smith, Eneko Villanueva, Maria Marti-Solano, Mie Monti, Mariavittoria Pizzinga, Dan-Mircea Mirea, Manasa Ramakrishna, Robert F. Harvey, Veronica Dezi, Gavin H. Thomas, Anne E. Willis, and Kathryn S. Lilley. Comprehensive identification of RNA-protein interactions in any organism using orthogonal organic phase separation (OOPS). Nature Biotechnology, 37(2):169–178, Feb 2019. ISSN 1546-1696. doi:10.1038/s41587-018-0001-2.

41. Benedikt M. Beckmann. RNA interactome capture in yeast. Methods (San Diego, Calif.), 118-119:82–92, Apr 2017. ISSN 1095-9130. doi:10.1016/j.ymeth.2016.12.008.

42. Thomas Conrad, Anne-Susann Albrecht, Veronica Rodrigues de Melo Costa, Sascha Sauer, David Meierhofer, and Ulf Andersson Ørom. Serial interactome capture of the human cell nucleus. Nature Communications, 7:11212, Apr 2016. ISSN 2041-1723. doi:10.1038/ncomms11212.

43. Francesco Itri, Daria M. Monti, Bartolomeo Della Ventura, Roberto Vinciguerra, Marco Chino, Felice Gesuele, Angelina Lombardi, Raffaele Velotta, Carlo Altucci, Leila Birolo, Renata Piccoli, and Angela Arciello. Femtosecond UV-laser pulses to unveil protein-protein interactions in living cells. Cellular and Molecular Life Sciences: CMLS, 73(3):637–648, Feb 2016. ISSN 1420-9071. doi:10.1007/s00018-015-2015-y.

44. Francesco Itri, Daria Maria Monti, Marco Chino, Roberto Vinciguerra, Carlo Altucci, Angela Lombardi, Renata Piccoli, Leila Birolo, and Angela Arciello. Identification of novel direct protein-protein interactions by irradiating living cells with femtosecond UV laser pulses. Biochemical and Biophysical Research Communications, 492(1):67–73, Oct 2017. ISSN 1090-2104. doi:10.1016/j.bbrc.2017.08.037.

45. Luke a Yates, Chris J Norbury, and Robert J C Gilbert. The long and short of microRNA. Cell, 153(3):516–9, apr 2013. ISSN 1097-4172. doi:10.1016/j.cell.2013.04.003.

46. Gunter Meister. Argonaute proteins: functional insights and emerging roles. Nature Reviews. Genetics, 14(7):447–459, Jul 2013. ISSN 1471-0064. doi:10.1038/nrg3462.

47. Pin Wang, Junfang Xu, Yujia Wang, and Xuetao Cao. An interferon-independent lncRNA promotes viral replication by modulating cellular metabolism. Science, 358(6366):1051–1055, Nov 2017. ISSN 1095-9203. doi:10.1126/science.aao0409.

48. Rastislav Horos, Magdalena Büscher, Rozemarijn Kleinendorst, Anne-Marie Alleaume, Abul K. Tarafder, Thomas Schwarzl, Dmytro Dziuba, Christian Tischer, Elisabeth M. Zielonka, Asli Adak, Alfredo Castello, Wolfgang Huber, Carsten Sachse, and Matthias W. Hentze. The Small Non-coding Vault RNA1-1 Acts as a Riboregulator of Autophagy. Cell, 176(5):1054–1067.e12, Feb 2019. ISSN 1097-4172. doi:10.1016/j.cell.2019.01.030.

