## Supplementary Figures for "Fast and unbiased purification of RNA-protein complexes after UV cross-linking"

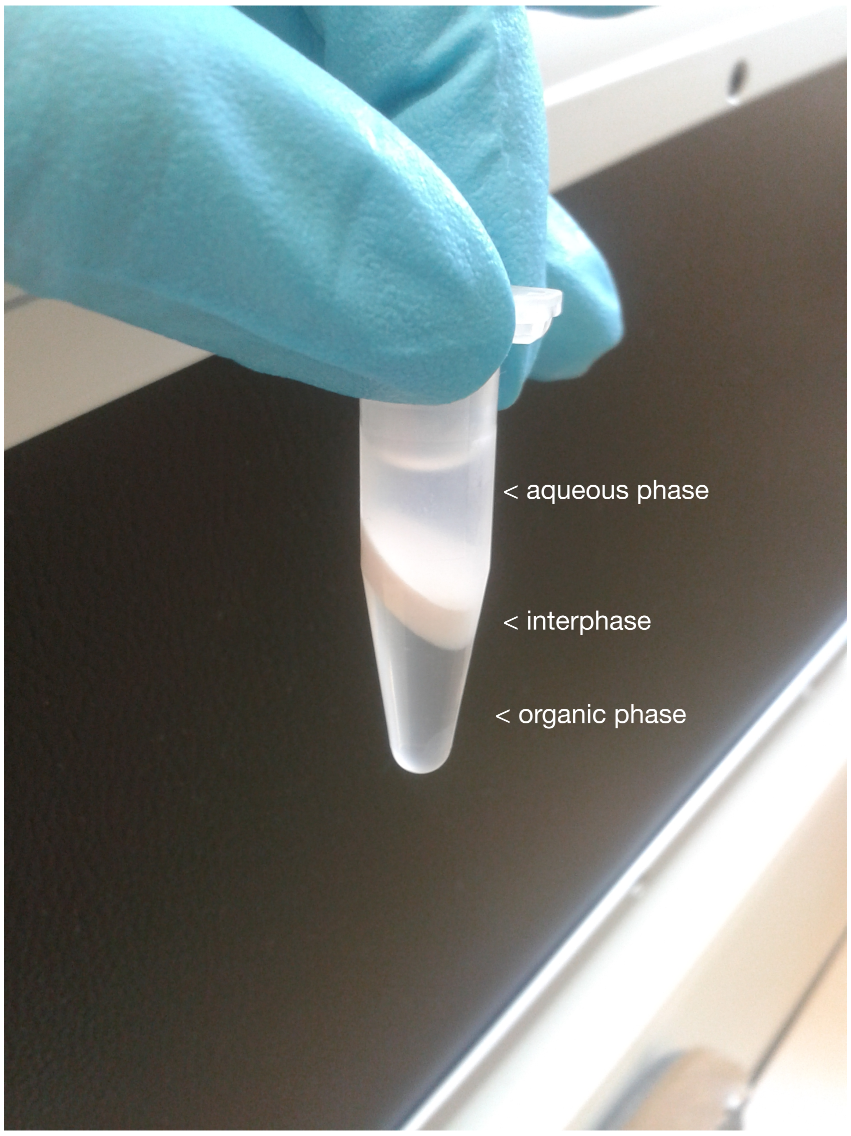


Supplementary Figure 1. PTex of HeLa cells (step 1). Notice the accumulation of cellular debris in the interphase and the turbidity of the aqueous phase, the latter serve as indicative of the presence of proteins and other soluble molecules.

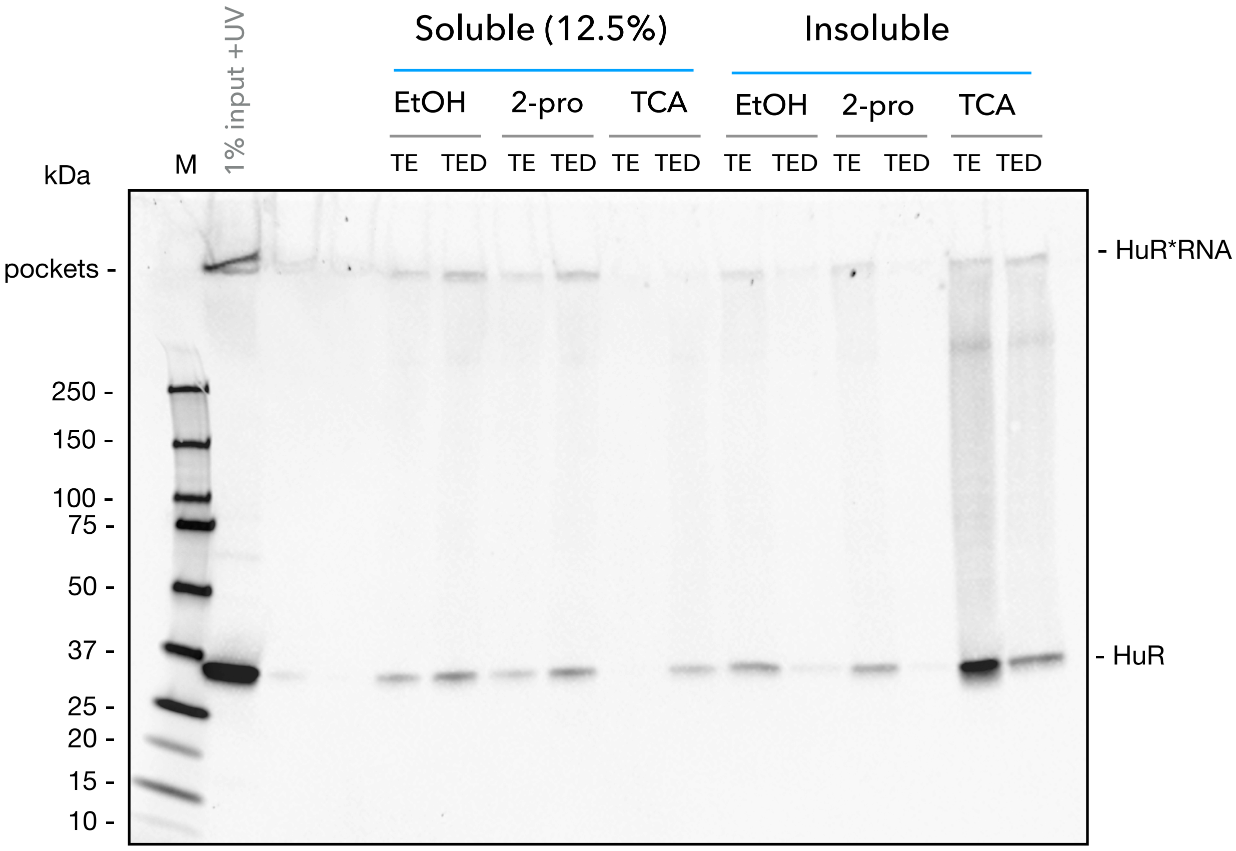


Supplementary Figure 2: Comparison of three protein precipitation methods: ethanol (EtOH), 2-propanol (2-pro) and trichloroacetic acid (TCA). HEK293 +UV lysates produced by PTex step 1, were precipitated with the indicated method (separated into soluble and insoluble fraction by centrifugation), electrophoresed and blotted to detect the protein HuR.

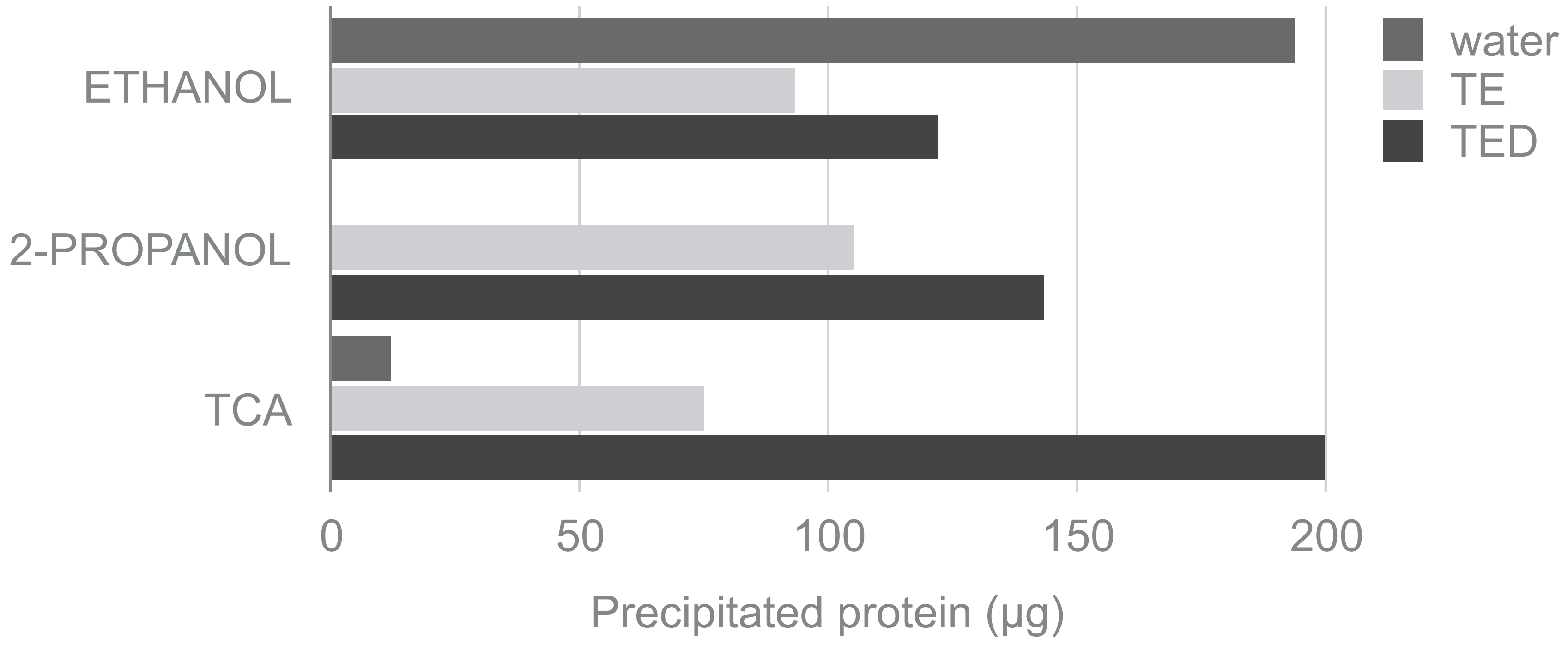


Supplementary Figure 3. Comparison of total protein (micrograms) obtained by three protein precipitation methods. HEK293 cell lysates (PTex, step 1) precipitated by ethanol, isopropanol or trichloroacetic acid. Pellets resuspended with 100 µl of water, TE or TED, incubated 20 min at 56 °C and centrifuged 1000 xg 2 min. Protein concentration determined as a function of the absorbance at 280 nm in a Nanodrop 2000. TE= 20 mM Tris, 1 mM EDTA, pH 7.8; TED= TE supplemented with 0.03% n-Dodecyl β-D-maltoside (DDM). Mean from two independent experiments.

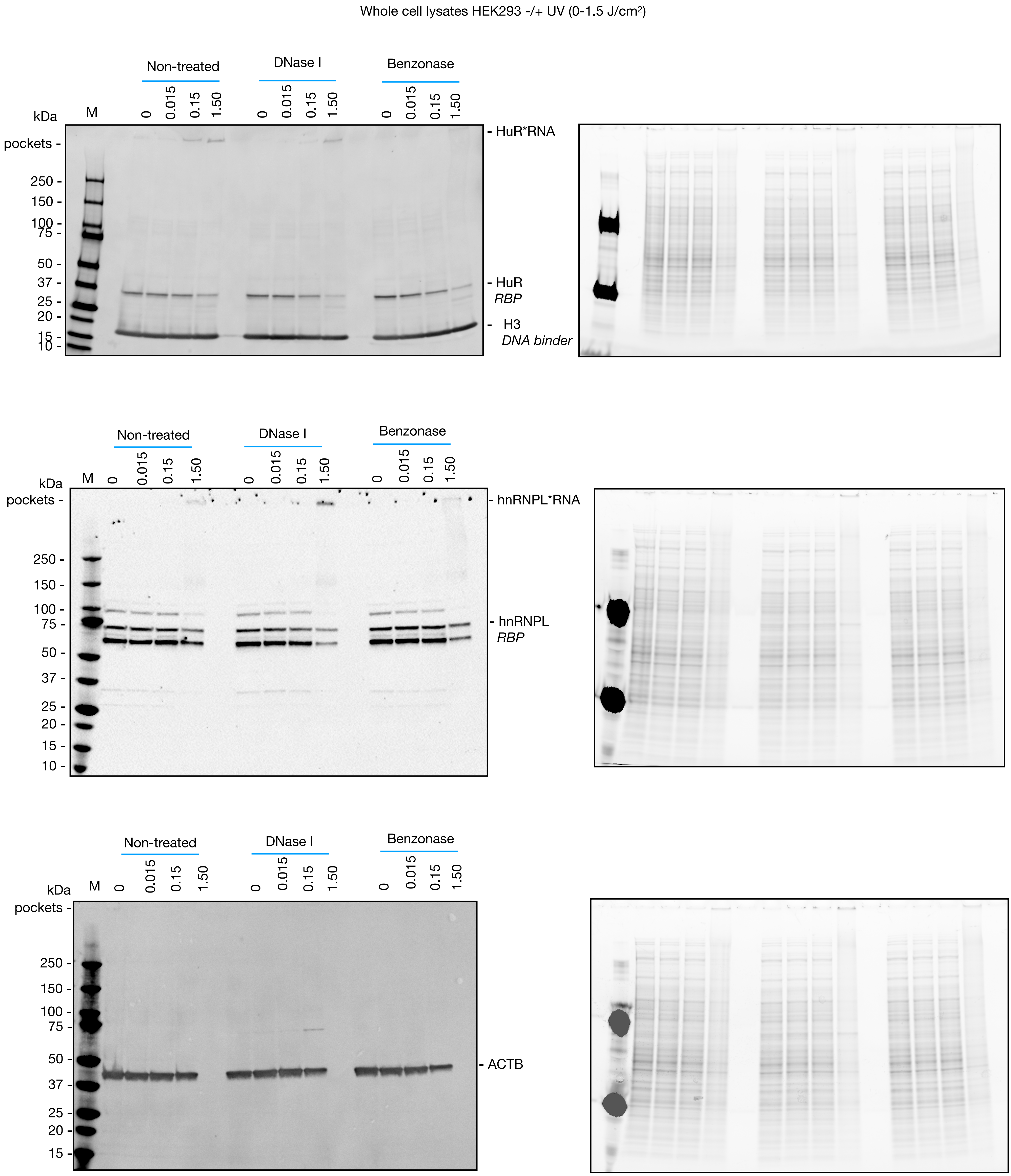


Supplementary Figure 4: blots for Fig. 2C. HEK293 cells were irradiated with an increasing dosage of $UV_{254\ nm}$, treated with DNAse I (0.2 U/µL) or Benzonase (25 U/µL). Lysed with Laemmli buffer, electrophoresed (right, Stain-Free visualisation system, BioRad) and blotted to reveal the indicated protein (left).

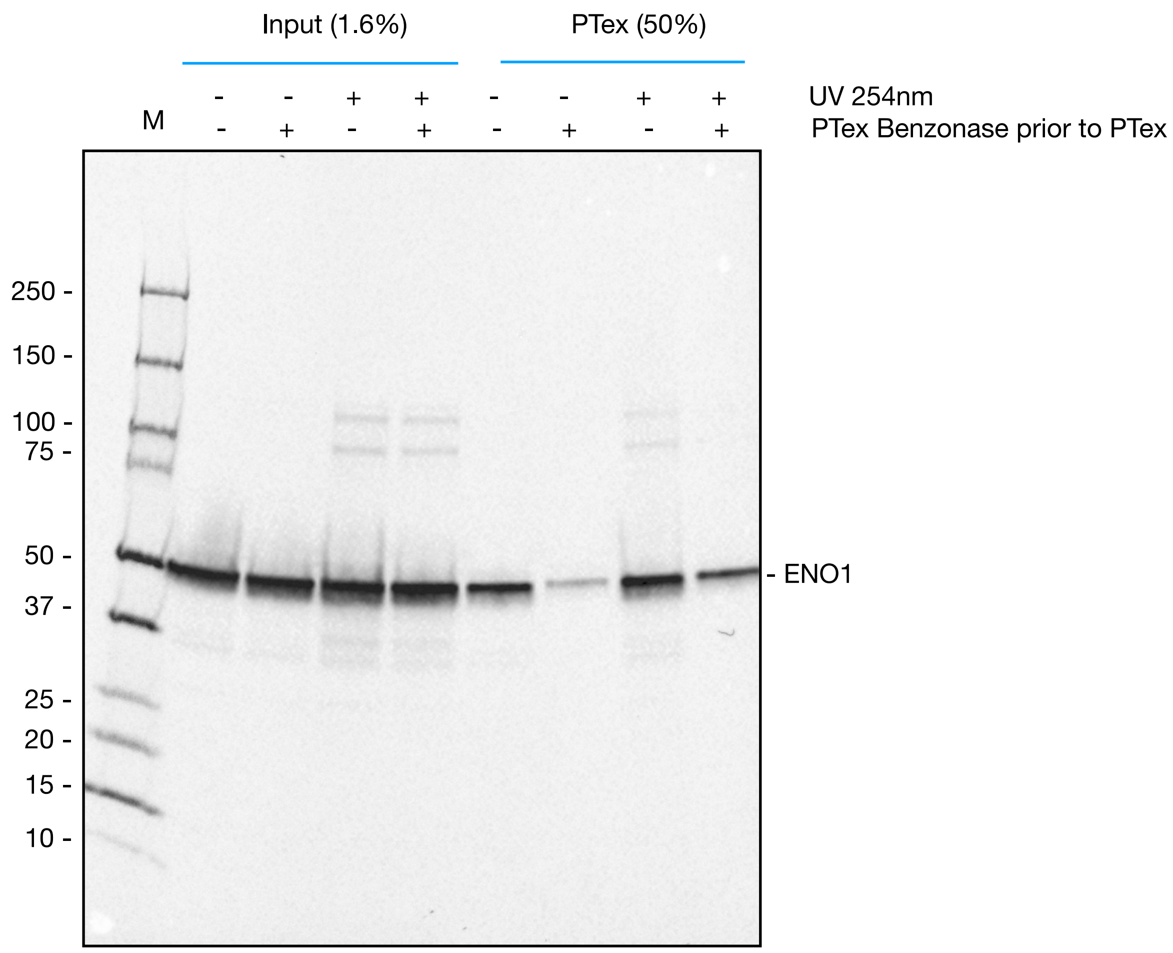


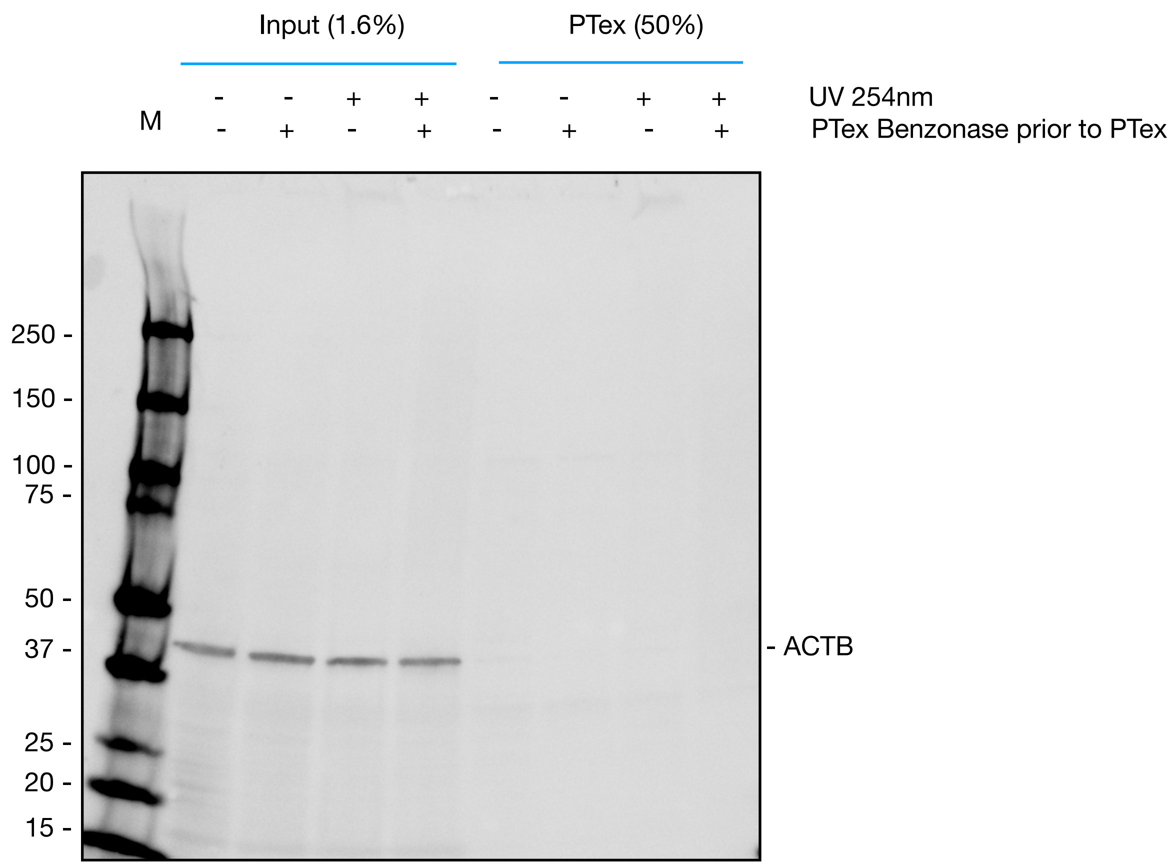


Supplementary Figure 5: blots for Figure 5B. HEK293 cellular suspensions (-/+ CL, 2-3x10$^6$ cells/mL) were incubated with 2 U/µL of benzonase at 37 °C, 1h, 1000 r.p.m. (Thermomixer, Eppendorf). Non-treated samples were kept as -Benzonase controls. For PTex, 600 µl per condition were used. Samples were electrophoresed and blotted to reveal the indicated protein.

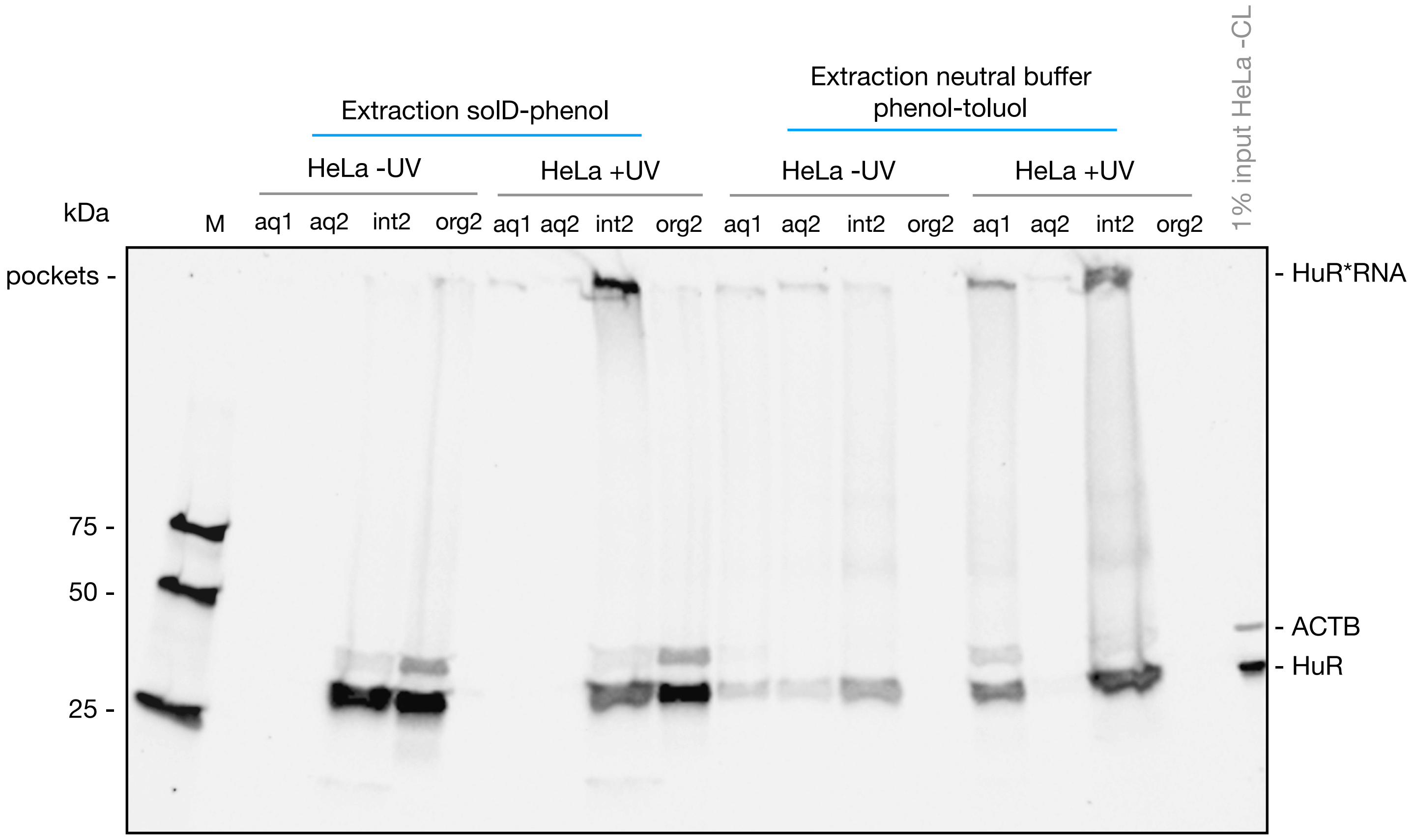


Supplementary Figure 6: Full blot for Fig. 3B, upper panel. Antibody against HuR (rabbit, 1:1000) and ACTB (mouse, 1:20.000).

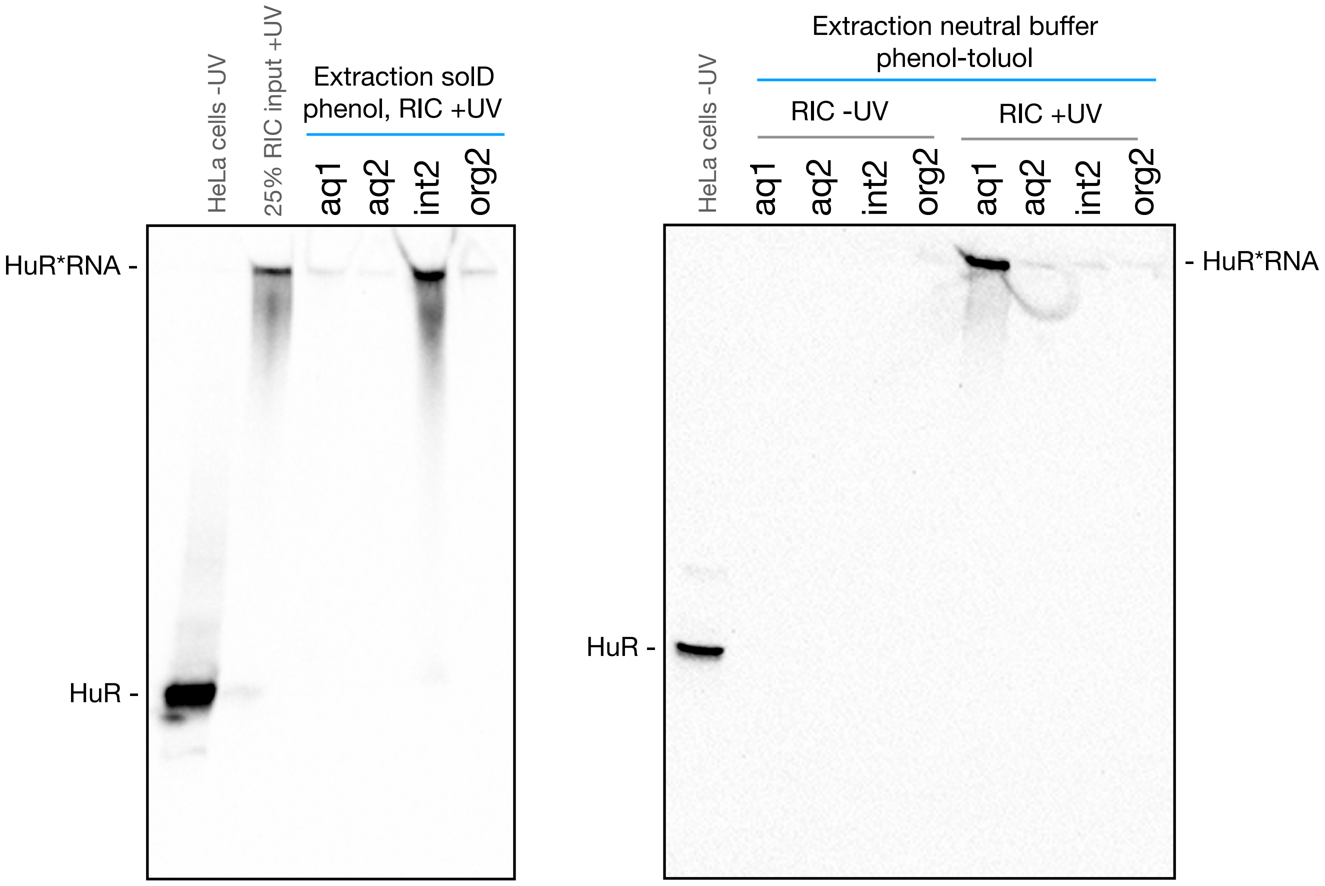


Supplementary Figure 7: Full blot for Fig. 3B, lower panel. Antibody against HuR (rabbit, 1:1000).

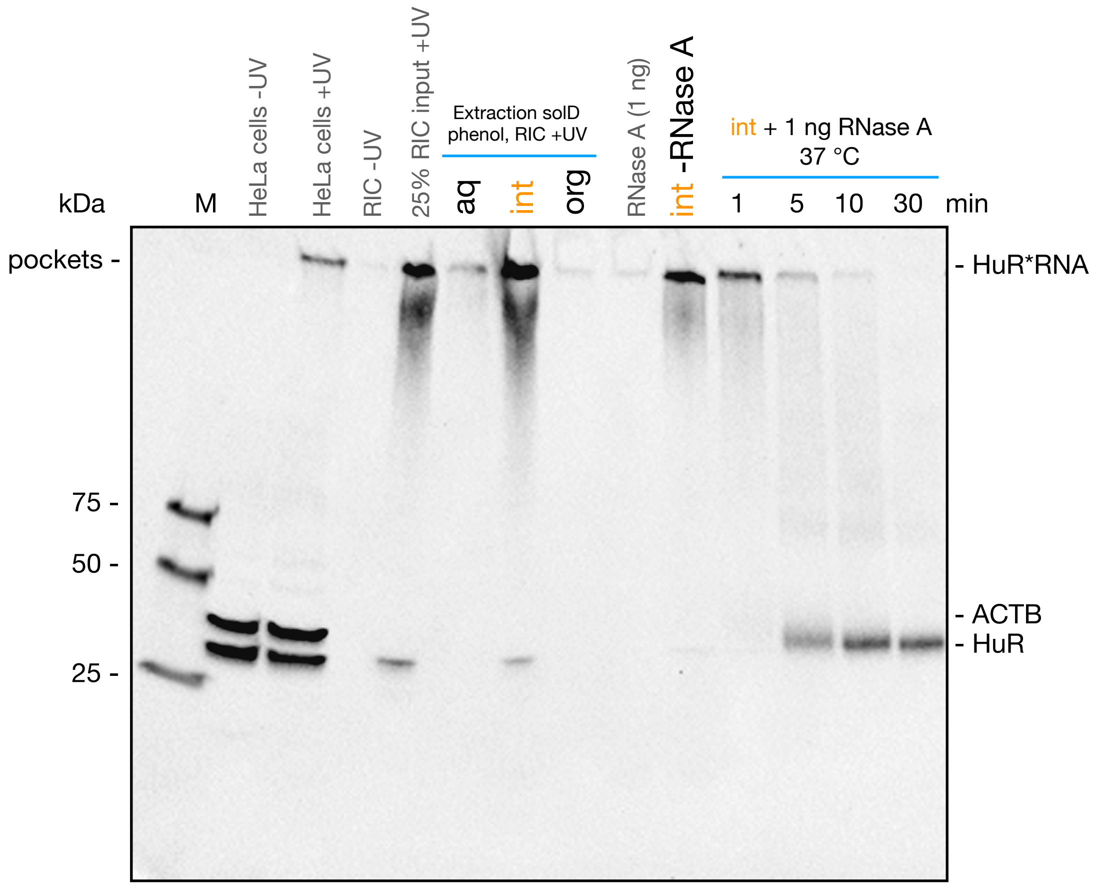


Supplementary Figure 8: Full blot for Fig. 3C

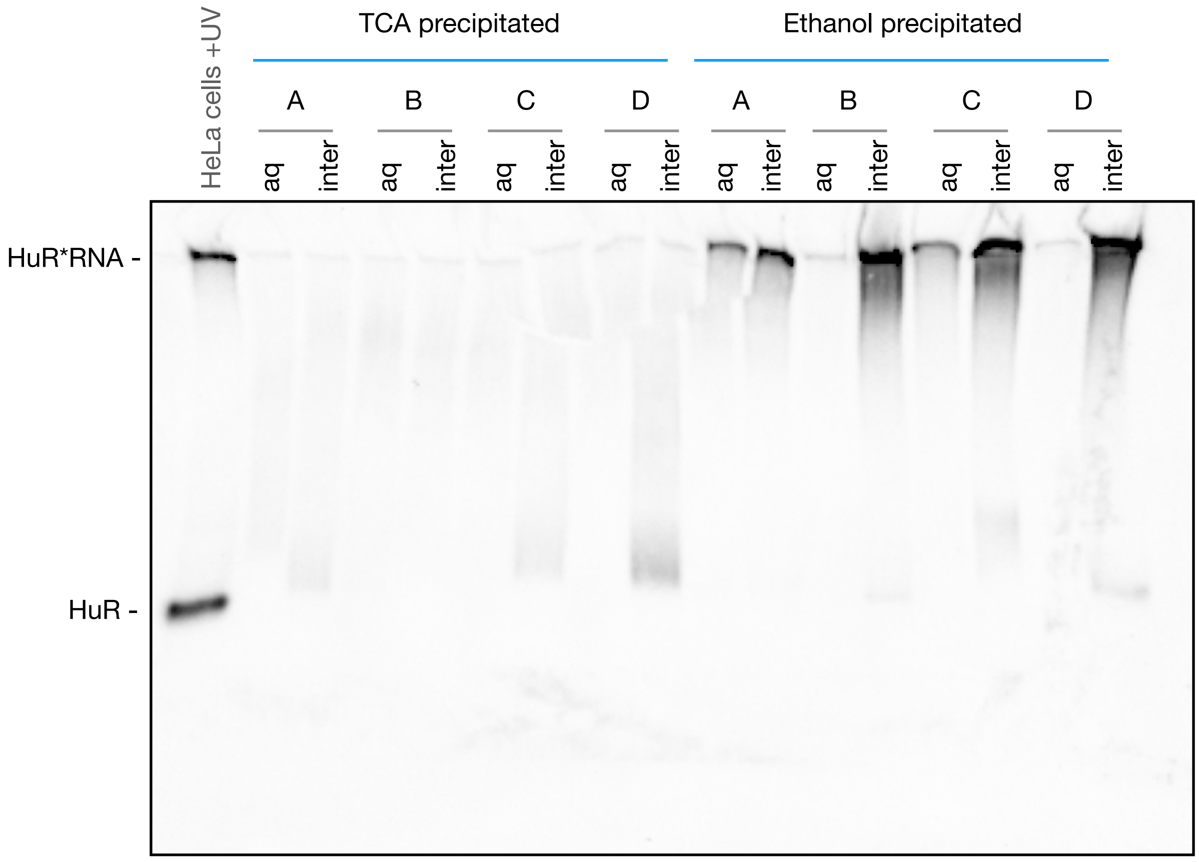


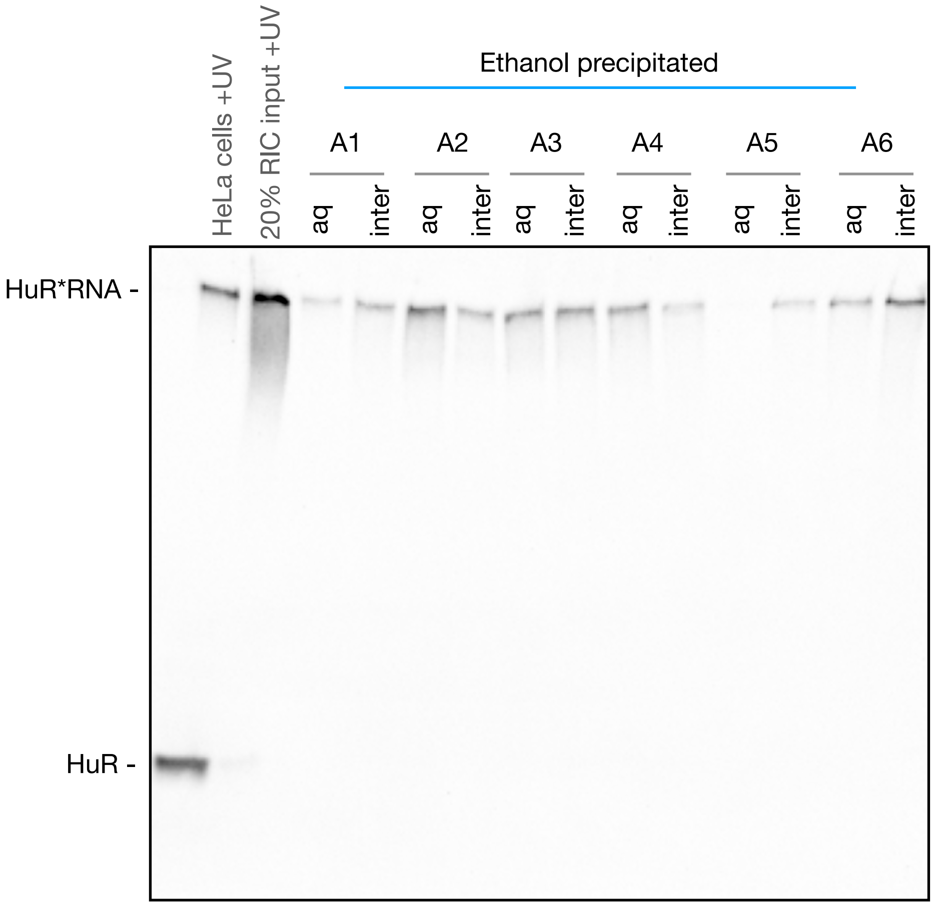


Supplementary Figure 9: Full blot for Fig. 3D.

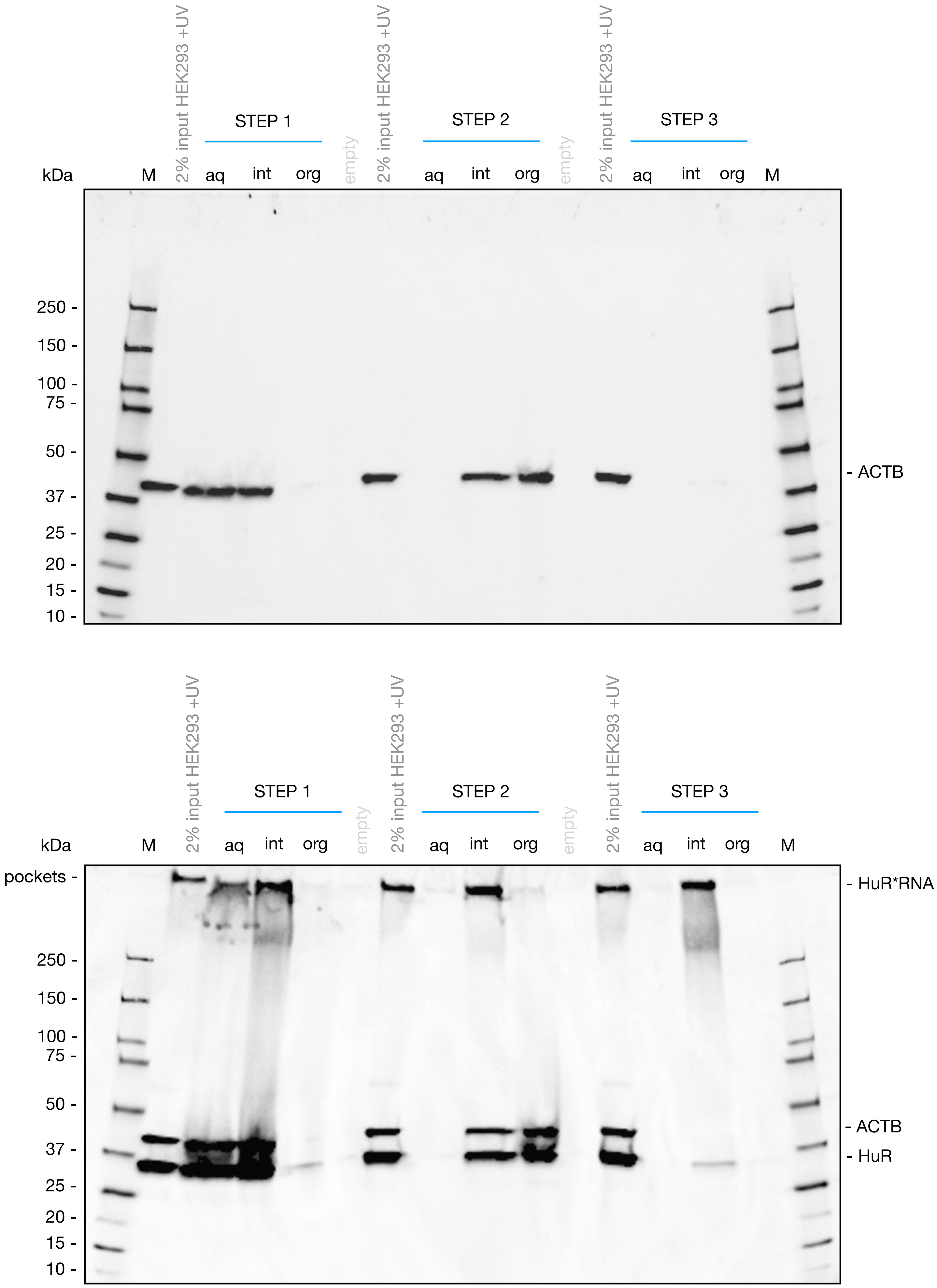


Supplementary Figure 10: Full blot for Fig. 4A. PTex intermediary steps. Membrane was firstly incubated with anti-ACTB (42 kDa, antibody 1:20.000; upper blot), developed and imaged; followed by a second blotting with anti-HuR (ELAV-1, 35 kDa, 1:1000; lower blot). Note that in interphase 1, only 20% of the material was used.

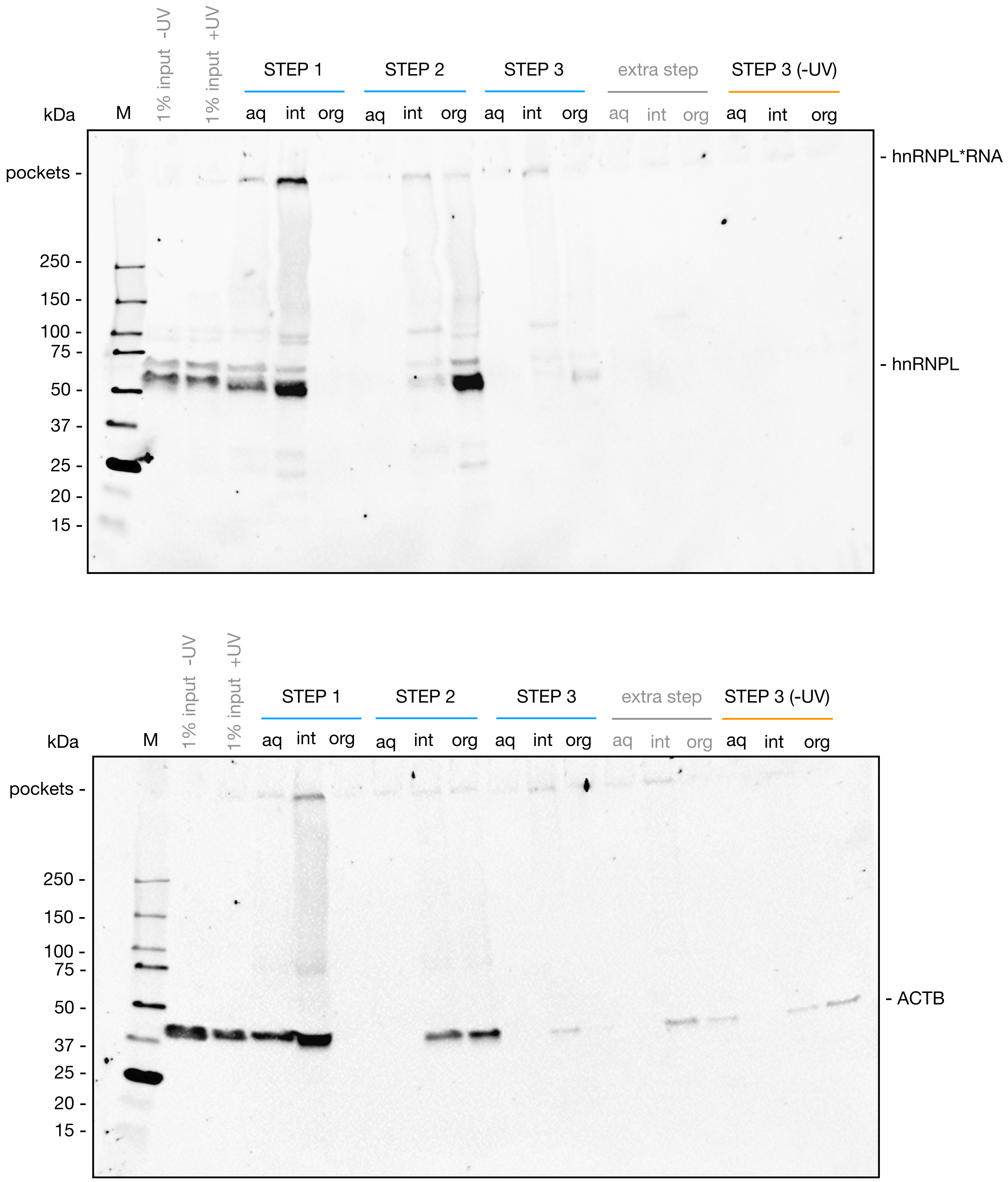


Supplementary Figure 11: full blots for figure 4B, PTex intermediary steps of HEK293 cells. The extra step consisted in the repetition of the step 2 at the end of the protocol aimed to further remove protein background, however, it resulted in a greater lost of the cross-linked fraction of the RBP while the non-RBP background remained unchanged. For the interphase 1, only 20% of the material was loaded.

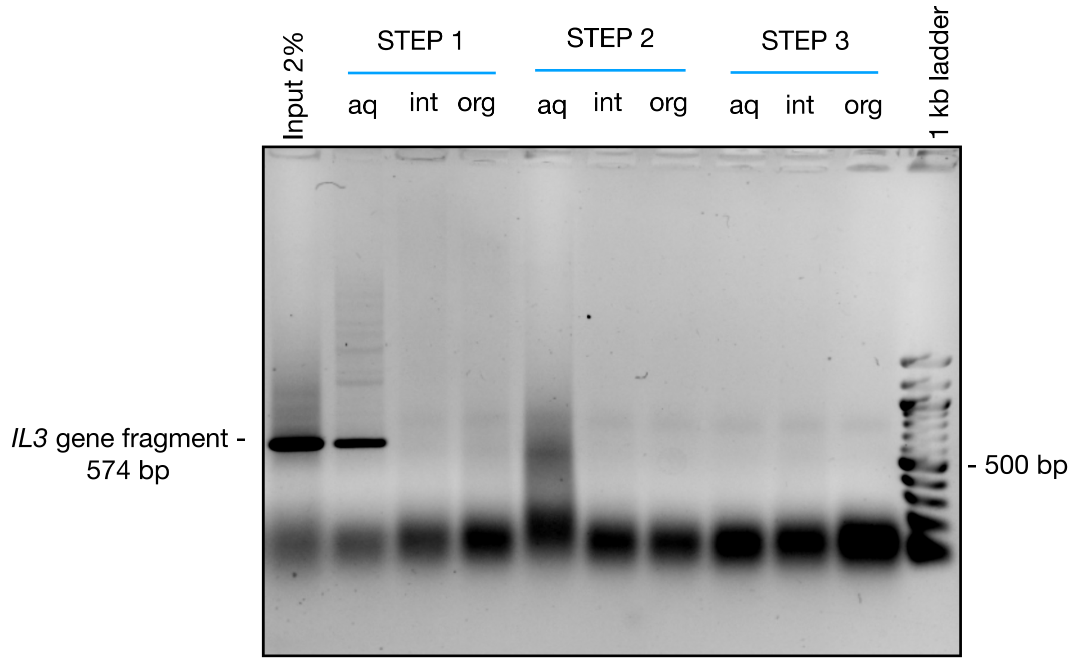


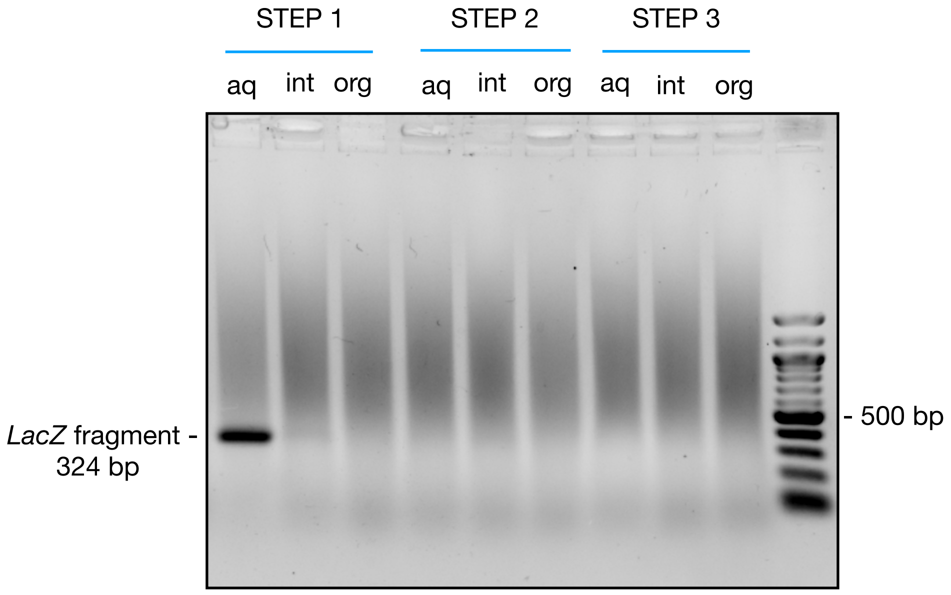


Supplementary Figure 12: PTex intermediary steps using pUC19 close plasmid or HEK293 genomic DNA as input. PCR was used to amplify the genes IL3 or LacZ.
