## Supplementary material for "Fast and unbiased purification of RNA-protein complexes after UV cross-linking": PTex Flyer

Purification of UV-crosslinked RNA-protein complexes by phenol-toluol extraction

Urdaneta & Beckmann / www.beckmannlab.de

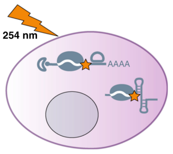


ALWAYS work under the hood and wear saftey gear.

$UV_{254nm}$ dosage must to be optimised in every case. Energies of 0.015-1.5 J/cm2 have been used for mammalian cells in culture.

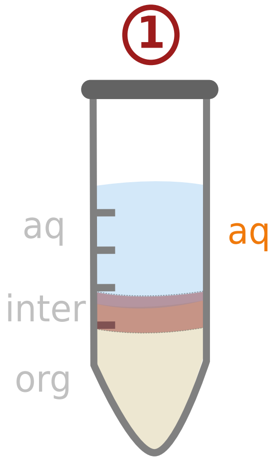


In a 2 mLtube (safe-cap) containing 2-6x$10^6$ cells in 0.6 mL PBS

Add:
0.2 mL neutral phenol
0.2 mL toluol
0.2 mL 1,3-bromo-chloro-propane (BCP)

Mix 1 min, RT in Thermomixer 2.000 r.p.m.
or Vortex max. speed.
Centrifuge 20.000 xg, 3 min, 4 °C.

Carefully transfer 0.4-0.5 mL of the resulting Aq to a new 2 mL tube containing 0.3 mL solution D, mix pipeting.

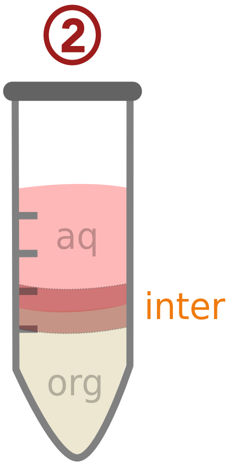


Add:
0.6 mL neutral phenol
0.2 mL BCP

Mix 1 min, RT in Thermomixer 2.000 r.p.m.
or Vortex max. speed.

Centrifuge 20.000 xg, 3 min, 4°C.

**Solution D:**
5.85 M guanidine isothiocyanate
31.1 mM sodium citrate.
25.6 mM N-lauryosyl-sarcosine
1% 2-mercaptoethanol

With a syringe and blunt needle (21G) remove 3/4th of the upper aq and 3/4th of the lower org. Keep the interphase in the same tube

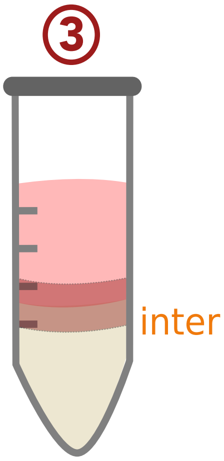


Add:
0.2 mL ethanol
0.4 mL H2O, mix briefly
0.4 mL neutral phenol
0.2 mL BCP

Mix 1min, RT in Thermomixer 2.000 r.p.m.
or Vortex max. speed.

Centrifuge 20.000 xg, 3 min, 4 °C.

With a new syringe remove 3/4th of the upper aq and 3/4th of the lower org.

Transfer the interphase to a 5 mL tube and add 9 vol. of ethanol. Incubate 30 min at 20 °C (or overnight).

Centrifuge at 20.000 xg, 30 min, 4 °C. Disolve the pellet in 30-50 µL $H_2O$ or Laemmli buffer.
